# Delving Below the Species Level To Characterize the Ecological Diversity within the Global Virome: An Exploration of West Nile Virus

**DOI:** 10.1101/2019.12.12.874214

**Authors:** Tong Kong, Kelly Mei, Ammie Wang, Danny Krizanc, Frederick M. Cohan

**Affiliations:** Departments of Mathematics & Computer Science, Wesleyan University, Middletown, Connecticut, USA; Departments of Biology, Wesleyan University, Middletown, Connecticut, USA

**Keywords:** Global Virome Project, host associations, ecotypes, genome sequence, microdiversity

## Abstract

Efforts to describe the diversity of viruses have largely focused on classifying viruses at the species level. However, substantial ecological diversity, both in virulence level and host range, is known within virus species. Here we demonstrate a proof of concept for easily discovering ecological diversity within a virus species taxon. We have focused on the West Nile Virus to take advantage of its broad host range in nature. We produced a genome-based phylogeny of world diversity of WNV and then used Ecotype Simulation 2 to hypothesize demarcation of genomes into 69 putative ecotypes (ecologically distinct populations), based only on clustering of genome sequences. Then we looked for evidence of ecological divergence among ecotypes based on differences in host bird associations within the Connecticut-New York region. Our results indicated significant heterogeneity among ecotypes for their associations with different bird hosts. Ecological diversity within other zoonotic viruses could be easily discovered using this approach. Opportunities for extending this line of research to human associations of virus ecotypes are limited by missing geographic metadata on human samples.

## Introduction

Recent virus spillovers have revealed our species’ vulnerability to infection from wild animals. Our domestic animals have recently conveyed several deadly bat viruses to humanity, including MERS (Drosten et al. 2014), Hendra (Plowright et al. 2011), and Nipah (Daszak et al. 2013); the bushmeat industry has brought us SARS (Yip et al. 2009); and human encroachment on wild lands has recurrently brought us Ebola (Rulli et al. 2017). Moreover, these infamous examples are not the whole story—on average, more than two new viruses are discovered in humans each year (Woolhouse et al. 2008). The undiscovered viruses residing in animal reservoirs are estimated to number 1.67 million, and about half of these are expected to have the capacity for infecting humans (Carroll et al. 2018).

Until recently, discovering the viruses that infect humanity from other animals has been a catch-up game. Starting when people get sick from a novel infection, virologists work urgently to characterize the virus, develop model systems, and produce vaccines and treatments. The Emerging Pandemic Threats project of USAID and the Global Virome Project have aimed to go beyond catch-up to *anticipate* future viral spillovers (Morse et al. 2012, Carroll et al. 2018). The Emerging Pandemic Threats project has aimed to identify the geographical regions, animal reservoirs, and human activities most likely to yield a future pandemic, and the GVP aims to discover, characterize, and sequence the genome of every virus on the planet that could possibly infect humans, all before the viruses can infect humanity. The GVP’s proposers have argued that if we limit our focus to the virus families most likely to infect humans, as well as the host taxa with the greatest frequency of yielding spillovers to humans, the project’s costs will in the end be much cheaper than paying for the damage from a pandemic (Carroll et al. 2018).

In its prospectus plan, the Global Virome Project focuses on finding new viruses that have important biological differences from already known species (Carroll et al. 2018). The GVP does not explicitly state the desired taxon rank at which new viruses should be sought, but it will likely aim to find viruses that are unique in their transmission, pathogenicity, cell and tissue tropism, host range, virion morphology, and antigenic relatedness—that is, viruses that meet the criteria of being distinct species taxa (Simmonds et al. 2017).

Virologists are increasingly discovering new virus species from molecular data, where a given level of sequence divergence suggests that an unknown virus is sufficiently distinct in its biological properties from other described species. For any given group of viruses, virologists have estimated a level of sequence divergence that correlates with significant differences in the properties of the viruses (Simmonds and Aiewsakun 2018). Recently, bioinformatic approaches have created algorithms to demarcate biologically significant virus groups on the basis of forming distinct sequence clusters (Lauber and Gorbalenya 2012, Yu et al. 2013, Bao et al. 2014). Sequence-based approaches hold promise for providing benchmark levels of sequence divergence that will allow discovery of biologically unique groups within a given virus group. The Global Virome Project may take advantage of these benchmarks in deciding whether an unknown virus is of sufficient interest.

Nevertheless, we cannot yet be certain that taxonomy of viruses at the species level is equipped to identify all the important ecological diversity among viruses. This is because extremely closely related viruses assigned to the same species taxon may be importantly different in their ecological features. Bacteriologists have long appreciated that a species taxon may contain important ecological diversity (Konstantinidis and Tiedje 2005, Denef et al. 2010, Shapiro et al. 2012, Cohan 2017), and significant diversification within species is becoming evident in the world of virology as well (Marston and Amrich 2009).

This is readily apparent in the case of bacteriophages. A recent study of microdiversity of marine phages revealed that extremely close relatives differed in the depths and bacterial hosts they were adapted to (Warwick-Dugdale et al. 2019). Likewise, animal viruses within a single virus species taxon have diversified to infect different hosts. For example, lineages within the Rabies Virus species taxon have diverged in the bat species they infect (Streicker et al. 2010). Moreover, Rabies lineages associated with different bats have shown a history of divergent selection in proteins involved in host cell interaction (Streicker et al. 2012).

Viruses within a single species have also diverged in their levels of virulence toward their animal hosts. For example, among subgroups of HIV-1 Group M, the C subgroup reproduces and causes disease in humans much less aggressively than other subgroups (Ariën et al. 2007, Blanquart et al. 2016). Within West Nile Virus, the focus of the present paper, major clades have been shown to differ in the viremia level caused in their bird hosts (Duggal et al. 2014), as well as in transmission from their mosquito vectors (Ebel et al. 2004, Hadfield et al. 2019). If viruses within a species taxon typically differ in either host specificities or virulence, then the search for new viruses may benefit from efforts to delve below the species level.

Here we present a test case for routine discovery of potentially novel pathogens within a virus species. We have focused on West Nile Virus, a zoonotic virus that recurrently spills over to humans from birds by way of mosquitoes. WNV is a broadly generalist virus—it infects hundreds of species of birds and is transmitted by scores of mosquito species (Colpitts et al. 2012, Reisen 2013). Our study takes advantage of this virus’s ecological generalism.

We have investigated whether phylogenetic clades within Cluster 4 of West Nile Virus, which has spread throughout the Americas (Mann et al. 2013), differ in their likelihood to be sampled from different bird species. Our approach was to hypothesize ecologically distinct populations (ecotypes) of West Nile Virus, using the algorithm Ecotype Simulation 2 (Koeppel et al. 2008, Francisco et al. 2014). The algorithm analyzes the sequence diversity of a rarely sexual or entirely asexual group to identify sequence clusters that are likely to be ecologically distinct (ecotypes), without using any ecological data. When ES2 has previously been applied to closely related bacteria coexisting within the same region, the putative ecotypes have been confirmed to be ecologically distinct through differences in habitat associations and physiology(Connor et al. 2010, Koeppel et al. 2013, Becraft et al. 2015). The algorithm should be appropriate to apply to West Nile Virus, since this virus recombines extremely rarely (Pickett and Lefkowitz 2009).

We will demonstrate that within one geographical region, putative ecotypes of West Nile Virus identified by Ecotype Simulation are quantitatively different in the host bird species they tend to infect, suggesting ecological diversification within the virus species. We will argue that a within-species focus may be valuable in discovering viruses with differing ecological niches.

## MATERIALS AND METHODS

### Aligning genomes

We obtained 1590 full-genome sequences of West Nile Virus from the NCBI Virus Variation database (Hatcher et al. 2017) in September 2018, drawing on the global diversity of WNV genome sequences from all mosquito and vertebrate hosts sampled from 55 regions on 6 continents. Identical sequences were collapsed, leaving 1319 unique sequences. The whole-genome sequences extracted varied in length from 10299 to 11355 bp. A built-in MUSCLE sequence alignment tool from the NCBI Virus Variation database constructed and exported an alignment file in FASTA format.

We used as an outgroup the whole genome sequence of Zika Virus strain KX893855, provided by NCBI. Zika and West Nile Virus are both in the *Flavivirus* genus, and Zika has a similar genome content and nucleotide length as WNV (Gupta et al. 2016, Rey et al. 2018). The Zika sequence was appended to the WNV sequence alignment through the “-add” function of MAFFT (Katoh and Standley 2013). We wrote a program, removeGaps.py, to remove gaps in the alignment file: https://github.com/tkong233/Global-Virome-Project. Applying the algorithm did not change the eventual demarcations of viruses into putative ecotypes. All programs written for this study are available through this Github link.

### Constructing the phylogenetic Tree

We used FastTree to construct a maximum-likelihood phylogeny from the genome alilgnment in Newick format, which was visualized by MEGA7. We then rooted the tree to the Zika virus outgroup within MEGA7’s Tree Explorer window.

### Demarcating putative ecotypes

We inputted both the alignment and Newick tree files into the Ecotype Simulation 2 algorithm. ES2 models the sequence diversity within an asexual or rarely-sexual clade to predict ecologically distinct populations from patterns of sequence clustering only, without using ecological data (Koeppel et al. 2008, Francisco et al. 2014) https://github.com/sandain/ecosim/releases. The log file of ES2 yielded ecotype demarcations, which were converted into an Excel file using our Python program dataFrameGen.py. The unique sequences within each ecotype are listed in Supplementary Table 1.

### Host associations of WNV putative ecotypes with taxonomic groupings of birds

One geographical region of particular interest comprised the adjacent states of Connecticut and New York in the USA, owing to sampling there of 362 birds providing West Nile Virus. Two bird species were abundantly sampled: the American crow (*Corvus brachyrhynchos*, 243 sequences), and the blue jay (*Cyanocitta cristata*, 80 sequences). Both these species are within the Corvidae family of the perching-bird order, Passeriformes. The remaining 39 birds sampled from Connecticut and New York included 20 other birds within the Passeriformes, 10 birds within the hawk-like order Accipitriformes, and 9 birds within the owl order Strigiformes. We tested for host association differences among putative ecotypes from a matrix of 5 taxonomic groups of birds (blue jays, American crows, other passeriforms, Accipitriformes, and Strigiformes) X 23 putative ecotypes, noting that putative ecotypes sampled fewer than 5 times from birds in Connecticut and New York were discarded from the analysis (Table 1). The matrix was analyzed by a contingency test of association using the G test in the DescTool package within RStudios. Finally, a phylogenetic tree containing only viruses sampled from Connecticut and New York birds was created with MAFFT, FastTree, and MEGA7 (Figure 1).

**Table 1.** Host associations of WNV putative ecotypes with five taxonomic groups of birds in the Connecticut-New York region. Listed are the total numbers of viruses from each ecotype infecting each host category, without regard to whether the sequences were unique. Included are those ecotypes with at least five viruses sampled from the Connecticut-New York region.

|  | American crows | Blue jays | Other<br>Passeriforms | Accipitriformes | Strigiformes |
| --- | --- | --- | --- | --- | --- |
| Ecotype 2 | 9 | 3 | 1 | 0 | 0 |
| Ecotype 3 | 10 | 0 | 0 | 0 | 0 |
| Ecotype 4 | 35 | 9 | 1 | 1 | 0 |
| Ecotype 15 | 3 | 3 | 0 | 0 | 1 |
| Ecotype 20 | 9 | 1 | 0 | 0 | 0 |
| Ecotype 21 | 5 | 2 | 0 | 0 | 0 |
| Ecotype 23 | 7 | 4 | 0 | 1 | 0 |
| Ecotype 24 | 7 | 0 | 3 | 1 | 0 |
| Ecotype 30 | 6 | 4 | 0 | 0 | 0 |
| Ecotype 35 | 3 | 6 | 3 | 0 | 0 |
| Ecotype 38 | 13 | 1 | 0 | 0 | 0 |
| Ecotype 39 | 4 | 6 | 0 | 0 | 0 |
| Ecotype 40 | 19 | 8 | 6 | 0 | 3 |
| Ecotype 41 | 9 | 2 | 1 | 0 | 0 |
| Ecotype 47 | 7 | 4 | 1 | 0 | 0 |
| Ecotype 48 | 11 | 5 | 0 | 0 | 0 |
| Ecotype 50 | 8 | 10 | 0 | 0 | 1 |
| Ecotype 51 | 8 | 2 | 1 | 1 | 0 |
| Ecotype 52 | 13 | 1 | 0 | 1 | 0 |
| Ecotype 53 | 8 | 2 | 0 | 1 | 2 |
| Ecotype 56 | 13 | 4 | 0 | 1 | 2 |
| Ecotype 58 | 11 | 2 | 1 | 0 | 0 |
| Ecotype 59 | 23 | 1 | 2 | 3 | 0 |

**Figure 1.**
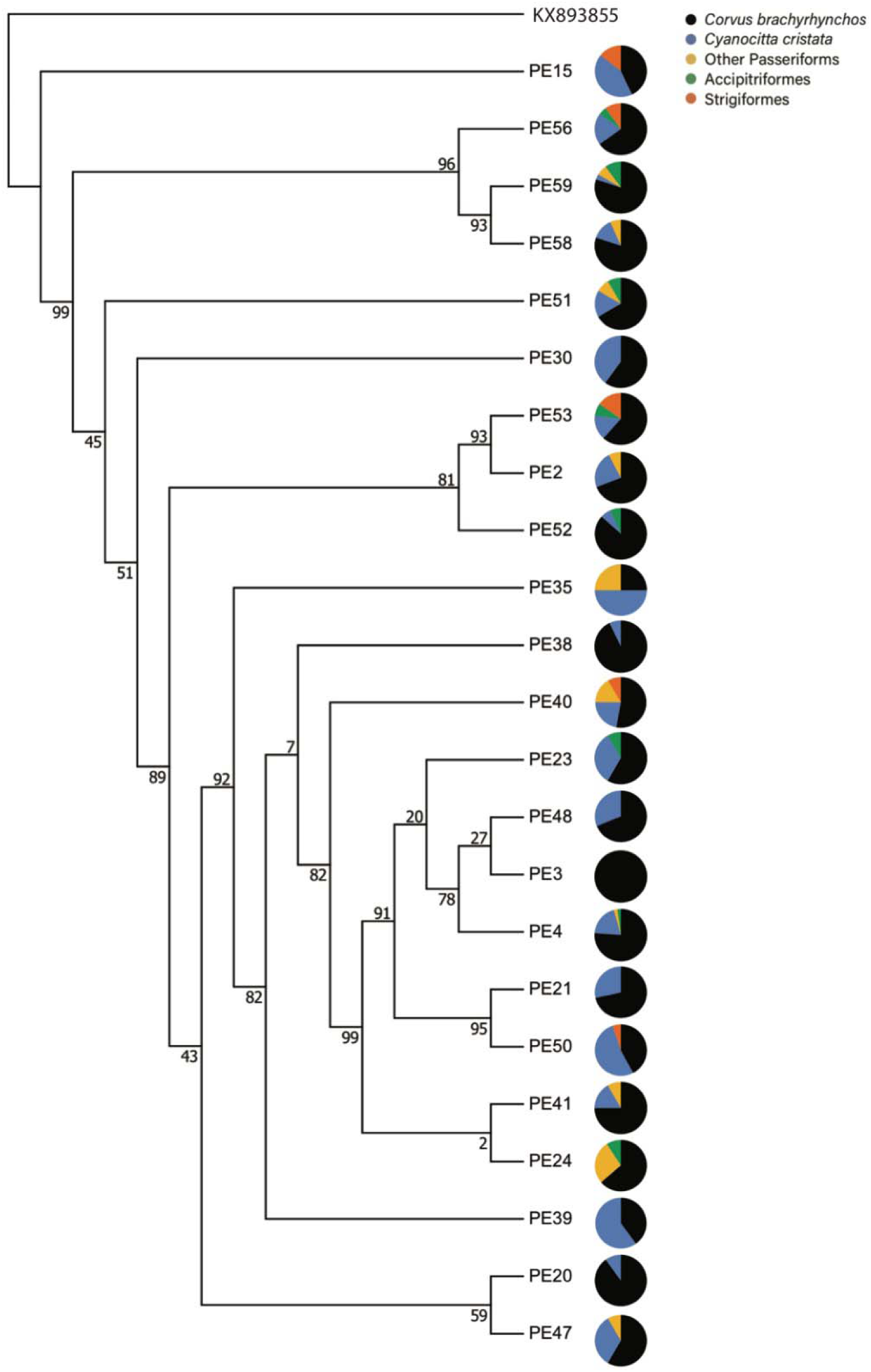
Host associations of WNV putative ecotypes with five taxonomic groups of birds in the Connecticut-New York region. The Zika virus strain KX893855 served as the outgroup.

## RESULTS

### Demarcation of WNV sequences into putative ecotypes

Based on the 1319 unique whole-genome sequences of WNV sampled worldwide, the Ecotype Simulation algorithm hypothesized and demarcated 69 putative ecotypes (Table 1, Supplementary Table 1).

### Associations of WNV putative ecotypes with bird taxa sampled in Connecticut and New York

From the Connecticut-New York region, we focused on WNV sequences from five taxonomic groups: the two bird species within the Passeriformes order that were most abundantly sampled (American crows and blue jays), various species less well sampled within the Passeriformes (“other passeriforms”), and two groups constituting the Accipitriformes and Strigiformes orders (Table 1). A contingency test yielded a highly significant heterogeneity among putative ecotypes for their associations with these host groups (G = 133.15, df = 84, P = 0.000514) (Figure 1). Among the most salient ecotype differences was the range of relative frequencies of American crows and blue jays, with for example, American crows ranging from 25% to 100% among the 23 putative ecotypes analyzed (Table 1).

## DISCUSSION

### Differences among WNV putative ecotypes in their associations with bird host species

Our aim was to identify important ecological and evolutionary divergence within a virus species taxon already known to be pathogenic for humans. We focused on West Nile Virus because it infects a variety of hosts and hence may have diversified by host specificity. We found that West Nile clades differed in their associations with bird hosts within the Connecticut-New York region. For example, the frequency of WNV samples in American crows varied from 25% to 100% among the 23 putative ecotypes analyzed. Because this study was circumscribed to one geographical region, where the bird hosts, mosquito vectors, and the viruses could disperse, the host associations are likely due to some intrinsic ecological difference (or differences) among the ecotypes.

One possible explanation for the host associations may be that some WNV lineages have evolved to reach higher blood titers in some bird species than in others, such that virologists are better able to detect them there. Alternatively, WNV lineages may have diversified by the mosquitoes they can most successfully infect (as seen for lineages within a fungal species) (Zebold et al. 1979). Provided that mosquito species differ in their likelihood to feed on one species of bird over another, virus diversification by mosquito vector could lead to the host associations we have observed.

### Limited opportunity to extend this analysis to associations of WNV lineages with humans

It is tempting to extend this line of analysis to compare putative ecotypes for their tendencies to associate with humans. Sufficient genome sequence data are available for this purpose, with WNV genome sequences from 569 birds and 69 humans (nearly all asymptomatic) sampled in the USA. Unfortunately, geographical metadata are entirely lacking for the human samples, beyond that they stem from the USA. The omission of the metadata is exacerbated by the fact that WNV overwinters throughout the continental USA (Reisen 2013) and so has developed some endemicity of lineages in geographic regions within the country (Hepp et al. 2018). Therefore, one could not absolutely ensure that any observed associations of putative ecotypes with humans did not emerge as an artifact of differences in where humans and birds were sampled.

Omission of important details in environmental metadata has repeatedly limited microbial ecologists’ ability to study the genetics of microbial adaptations to environments and hosts (Cohan 2012). We suggest that virus genome sequences should report at least minimal geographical and environmental metadata about where each sample was collected, as guided by Minimum Information about any(x) Sequence (MIxS) (Yilmaz et al. 2011).

Nevertheless, we tentatively explored differences in our WNV putative ecotypes in their tendencies to be sampled from humans versus birds. To try to reduce geographical bias, we focused only on ecotypes that were present in regions where both humans and birds were sampled (that is, ecotypes containing at least one bird and one human sample). The 18 putative ecotypes meeting this criterion were considerably different in their frequencies of human samples, ranging from 4.8% to 65%, with a highly significant heterogeneity in a contingency test (G = 76.828, df = 17, P = 1.40e-09) (Supplementary Table 2).

One possible explanation may be that some virus ecotypes reproduce more aggressively within humans, leading to a greater probability of being detected in blood samples in asymptomatic humans. Alternatively, some ecotypes may reproduce more successfully within the peridomestic mosquitos that most frequently bite humans, such as *Aedes albopictus*. This would lead to greater rates of infection of these ecotypes in humans. Finally, and unavoidably with the present data set, the ecotype differences could reflect a geographical bias in sampling humans.

### Potential implications of WNV bird associations for human spillovers

Evolutionary diversification of WNV ecotypes to different bird species could possibly impact the likelihood of spillover to humans. This is because some animal species serve better as an “ecological bridge” to transmit viruses to humans. For example, humans would never have become infected by Hendra Virus directly from its reservoir hosts in the *Pteropus* fruit bat genus. However, when the bats transmit the infection to horses, the virus is sufficiently amplified in horses’ saliva to infect the horses’ caretakers (NSW Government 2017). Also, pigs have served as an ecological bridge for infection of pigs’ caretakers by Nipah virus, again from *Pteropus* bats (National Center for Emerging and Zoonotic Infectious Diseases 2017). We suggest that evolution of a WNV ecotype that is particularly good at infecting a suburban backyard bird at high blood titers may result more frequently in human infections.

In some cases, a virus can more easily evolve adaptation to a new host species if it has previously evolved to infect an “evolutionary bridge” host. This is seen recurrently in Influenza, where pigs serve as a vessel for bird-adapted influenza to become adapted to mammals (Webster et al. 1992). Also, the Parvovirus lineage that eventually became Canine Parvovirus II is thought to have utilized its brief time infecting raccoons in evolving from a cat-infecting to a dog-infecting virus (Allison et al. 2012). Likewise, some bird species may possibly serve as an evolutionary bridge for WNV to more successfully infect humans, facilitating spillover by some WNV ecotypes more than others.

### Implications for the Global Virome Project

The GVP has generally sought to look for novel viruses that are outside of existing species taxa. However, virologists have found occasional evidence of important ecological divergence within virus species taxa (Ebel et al. 2004, Ariën et al. 2007, Streicker et al. 2010, Blanquart et al. 2016). Our present study presents a proof of concept that we can easily hypothesize putative ecotypes within a virus species taxon from genome sequence data, and then test the putative ecotypes for ecological distinctness by exploring for differences in host associations.

## Acknowledgements

We are grateful for funding from Wesleyan University, the John and Rosemarie Dooley Foundation, and from the Huffington Foundation.

## Data Availability Statement

The data we have created are available in Table 1 and Supplementary Tables 1 and 2.

## Ethical Statement

An ethical statement is not applicable as the data we have analyzed were collected from the NCBI database.

## SUPPORTING INFORMATION

**Supplementary Table 1. Sequence names of the 1319 unique WNV whole-genome sequences, as demarcated into putative ecotypes by Ecotype Simulation 2**.

**Supplementary Table 2. Host associations of WNV putative ecotypes with birds versus humans in the USA**. Listed are the total numbers of viruses from each ecotype infecting each host category, without regard to whether the sequences were unique. Included are those ecotypes with at least one human and one bird sample from the USA.

