## Supplementary Tables for "Delving Below the Species Level To Characterize the Ecological Diversity within the Global Virome: An Exploration of West Nile Virus"

**Supplementary Table 1. Sequence names of the 1319 unique WNV whole-genome sequences, as demarcated into putative ecotypes by Ecotype Simulation 2.**

Ecotype 1: [['FJ159131', 'Uranotaenia', 'Russia'], ['AY277251', 'Dermacentor marginatus', 'Russia'], ['FJ159129', 'Uranotaenia', 'Russia'], ['FJ159130', 'Uranotaenia', 'Russia']]

Ecotype 2: [['HM488250', 'Corvus brachyrhynchos', 'USA'], ['HM488183', 'Cyanocitta cristata', 'USA'], ['HQ671705', 'Culiseta melanura', 'USA'], ['KX547583', 'Corvus brachyrhynchos', 'USA'], ['AY712947', 'Aves', 'USA'], ['KX547542', 'Corvus brachyrhynchos', 'USA'], ['HQ671701', 'Culex salinarius', 'USA'], ['KX547480', 'Culex', 'USA'], ['KX547566', 'Culex', 'USA'], ['KJ501413', 'Corvus brachyrhynchos', 'USA'], ['KX547383', 'Culex', 'USA'], ['KX547570', 'Cyanocitta cristata', 'USA'], ['KX547400', 'Passer domesticus', 'USA'], ['KX547413', 'Corvus brachyrhynchos', 'USA'], ['KJ501495', 'Cyanocitta cristata', 'USA'], ['JQ700437', 'Culex pipiens', 'USA'], ['KX547437', 'Culex', 'USA'], ['KX547581', 'Culex', 'USA'], ['HQ671723', 'Corvus brachyrhynchos', 'USA'], ['DQ164186', 'Corvus brachyrhynchos', 'USA'], ['KX547591', 'Cyanocitta cristata', 'USA'], ['KX547338', 'Corvus brachyrhynchos', 'USA'], ['KX547469', 'Sciurus carolinensis', 'USA'], ['HM488174', 'Culex salinarius', 'USA'], ['DQ164190', 'Corvus brachyrhynchos', 'USA'], ['KX547205', 'Culex', 'USA'], ['KJ501479', 'Passer domesticus', 'USA']]

Ecotype 3: [['KX547199', 'Culex', 'USA'], ['KX547313', 'Corvus brachyrhynchos', 'USA'], ['HQ671729', 'Corvus brachyrhynchos', 'USA'], ['KJ501243', 'Corvus brachyrhynchos', 'USA'], ['KJ501244', 'Corvus brachyrhynchos', 'USA'], ['KJ501439', 'Corvus brachyrhynchos', 'USA'], ['KJ501215', 'Culex pipiens', 'USA'], ['KJ501246', 'Corvus brachyrhynchos', 'USA'], ['KJ501245', 'Corvus brachyrhynchos', 'USA'], ['KJ501248', 'Corvus brachyrhynchos', 'USA'], ['KJ501247', 'Corvus brachyrhynchos', 'USA'], ['KJ501249', 'Corvus brachyrhynchos', 'USA']]

Ecotype 4: [['KX547340', 'Culex', 'USA'], ['HM488169', 'Culex pipiens', 'USA'], ['HM488168', 'Culex pipiens', 'USA'], ['HM488166', 'Culex pipiens', 'USA'], ['HM488170', 'Culex pipiens', 'USA'], ['HM488167', 'Culex pipiens', 'USA'], ['JF920758', 'Culiseta melanura', 'USA'], ['KX547319', 'Corvus brachyrhynchos', 'USA'], ['KJ501505', 'Corvus brachyrhynchos', 'USA'], ['KJ501448', 'Corvus brachyrhynchos', 'USA'], ['KJ501517', 'Corvus brachyrhynchos', 'USA'], ['HM756659', 'Culiseta melanura', 'USA'], ['KX547405', 'Corvus brachyrhynchos', 'USA'], ['KJ501483', 'Cyanocitta cristata', 'USA'], ['KX547617', 'Culex', 'USA'], ['HM488231', 'Culiseta melanura', 'USA'], ['HM488232', 'Culiseta melanura', 'USA'], ['HM488222', 'Culiseta melanura', 'USA'], ['KJ501294', 'Corvus brachyrhynchos', 'USA'], ['KX547415', 'Corvus brachyrhynchos', 'USA'], ['KX547587', 'Turdus migratorius', 'USA'], ['KX547350', 'Culex', 'USA'], ['KX547616', 'Corvus brachyrhynchos', 'USA'], ['KX547329', 'Cyanocitta cristata', 'USA'], ['KX547382', 'Cyanocitta cristata', 'USA'], ['KX547516', 'Corvus brachyrhynchos', 'USA'], ['KX547342', 'Culex', 'USA'], ['KX547209', 'Corvus brachyrhynchos', 'USA'], ['KX547435', 'Corvus brachyrhynchos', 'USA'], ['KX547385', 'Cyanocitta cristata', 'USA'], ['JF920734', 'Culex restuans', 'USA'], ['KX547224', 'Culex', 'USA'], ['KX547433', 'Zenaida macroura', 'USA'], ['KJ501274', 'Corvus brachyrhynchos', 'USA'], ['HM488209', 'Ochlerotatus sticticus', 'USA'], ['KX547299', 'Corvus brachyrhynchos', 'USA'], ['KX547474', 'Sciurus carolinensis', 'USA'], ['KJ501496', 'Cyanocitta cristata', 'USA'], ['KJ501401', 'Corvus brachyrhynchos', 'USA'], ['KJ501263', 'Cyanocitta cristata', 'USA'], ['KX547444', 'Culex', 'USA'], ['KJ501291', 'Corvus brachyrhynchos', 'USA'], ['KJ501491', 'Corvus brachyrhynchos', 'USA'], ['KJ501306', 'Buteo jamaicensis', 'USA'], ['KX547398', 'Corvus brachyrhynchos', 'USA'], ['JF920752', 'Culex pipiens', 'USA'], ['KX547326', 'Culex', 'USA'], ['KX547359', 'Corvus brachyrhynchos', 'USA'], ['KX547250', 'Corvus brachyrhynchos', 'USA'], ['KF704158', 'Culex quinquefasciatus', 'USA'], ['JQ700440', 'Homo sapiens', 'USA'], ['KM012170', 'Homo sapiens', 'USA'], ['KM012174', 'Homo sapiens', 'USA'], ['MG004539', 'Culex quinquefasciatus', 'USA'], ['MG004529', 'Culex quinquefasciatus', 'USA'], ['KJ145827', 'Corvus brachyrhynchos', 'USA'], ['KX547600', 'Corvus brachyrhynchos', 'USA'], ['KJ501383', 'Cyanocitta cristata', 'USA'], ['KX547237', 'Corvus brachyrhynchos', 'USA'], ['HM756671', 'Corvus brachyrhynchos', 'USA'], ['KX547281', 'Corvus brachyrhynchos', 'USA'], ['KJ501359', 'Corvus brachyrhynchos', 'USA'], ['HM488219', 'Culex pipiens', 'USA'], ['KJ501305', 'Corvus brachyrhynchos', 'USA'], ['HM488140', 'Aedes vexans', 'USA'], ['KX547317', 'Corvus brachyrhynchos', 'USA'], ['HM756672', 'Corvus brachyrhynchos', 'USA'], ['KX547470', 'Corvus brachyrhynchos', 'USA'], ['KX547223', 'Corvus brachyrhynchos', 'USA'], ['KJ501371', 'Cyanocitta cristata', 'USA'], ['KX547374', 'Culex', 'USA'], ['KX547165', 'Cyanocitta cristata', 'USA'], ['HQ671724', 'Corvus brachyrhynchos', 'USA'], ['KJ501211', 'Culex tarsalis', 'USA'], ['JF415918', 'Cyanocitta cristata', 'USA'], ['KX547285', 'Corvus brachyrhynchos', 'USA'], ['KX547298', 'Culex', 'USA'], ['HM488214', 'Culex pipiens', 'USA'], ['KX547439', 'Corvus brachyrhynchos', 'USA'], ['HM756666', 'Corvus brachyrhynchos', 'USA'], ['HM488216', 'Culiseta melanura', 'USA'], ['KX547230', 'Culex', 'USA'], ['KX547371', 'Corvus brachyrhynchos', 'USA'], ['HM756678', 'Corvus brachyrhynchos', 'USA']]

Ecotype 5: [['KJ501446', 'Bubo virginianus', 'USA'], ['KJ501338', 'Cyanocitta cristata', 'USA'], ['KJ501400', 'Passer domesticus', 'USA'], ['KJ501218', 'Culex pipiens', 'USA'], ['KX547499', 'Corvus brachyrhynchos', 'USA'], ['JF488094', 'Corvus brachyrhynchos', 'USA'], ['HM488186', 'Corvus brachyrhynchos', 'USA'], ['JF488088', 'Culex pipiens', 'USA'], ['DQ080061', 'Cardinalis', 'USA'], ['KX547411', 'Culex', 'USA']]

Ecotype 6: [['KX547533', 'Corvus brachyrhynchos', 'USA'], ['DQ431703', 'Homo sapiens', 'USA'], ['DQ431701', 'Homo sapiens', 'USA']]

Ecotype 7: [['KX547229', 'Cyanocitta cristata', 'USA'], ['HM756670', 'Corvus brachyrhynchos', 'USA'], ['KX547573', 'Culex', 'USA']]

Ecotype 8: [['KX547438', 'Culex', 'USA'], ['JF920753', 'Culex restuans', 'USA'], ['KX547243', 'Corvus brachyrhynchos', 'USA']]

Ecotype 9: [['HM488192', 'Corvus brachyrhynchos', 'USA']]

Ecotype 10: [['KX547234', 'Culex', 'USA'], ['KX547328', 'Culex', 'USA']]

Ecotype 11: [['JF920731', 'Culex pipiens', 'USA'], ['KX547607', 'Culex', 'USA'], ['KX547562', 'Culex', 'USA'], ['KX547574', 'Culex', 'USA'], ['KX547417', 'Culex', 'USA'], ['KJ501538', 'Culicidae', 'USA'], ['KX547216', 'Culex', 'USA'], ['KX547380', 'Culex', 'USA'], ['KX547368', 'Culex', 'USA'], ['KJ501539', 'Culicidae', 'USA'], ['KX547464', 'Culex', 'USA'], ['KX547598', 'Corvus brachyrhynchos', 'USA'], ['KJ501534', 'Culicidae', 'USA'], ['KX547369', 'Culex', 'USA'], ['JQ700442', 'Homo sapiens', 'USA'], ['KX547495', 'Culex', 'USA'], ['KX547242', 'Culex', 'USA']]

Ecotype 12: [['KX547195', 'Corvus brachyrhynchos', 'USA'], ['KX547431', 'Culex', 'USA'], ['KX547295', 'Culex', 'USA'], ['HM488206', 'Corvus brachyrhynchos', 'USA'], ['KJ501536', 'Culicidae', 'USA'], ['KX547537', 'Culex', 'USA'], ['KJ501535', 'Culicidae', 'USA'], ['KX547279', 'Culex', 'USA']]

Ecotype 13: [['HM488238', 'Corvus brachyrhynchos', 'USA']]

Ecotype 14: [['KX547572', 'Culex', 'USA'], ['KY229068', 'Culex pipiens', 'USA'], ['KY229071', 'Culex pipiens', 'USA'], ['KY216150', 'Culex pipiens', 'USA'], ['KX547575', 'Culex', 'USA'], ['KY216149', 'Culex pipiens', 'USA'], ['KX547445', 'Culex', 'USA'], ['KX547559', 'Culex', 'USA'], ['KX547497', 'Culex', 'USA'], ['KJ501537', 'Culicidae', 'USA'], ['KJ501531', 'Culicidae', 'USA'], ['KJ501532', 'Culicidae', 'USA'], ['KX547214', 'Corvus brachyrhynchos', 'USA'], ['KM012185', 'Homo sapiens', 'USA'], ['KM012182', 'Homo sapiens', 'USA'], ['KJ501224', 'Culex tarsalis', 'USA'], ['KJ501229', 'Culex tarsalis', 'USA'], ['KJ501228', 'Culex tarsalis', 'USA'], ['KR868734', 'Culex tarsalis', 'USA'], ['KC333385', 'Cyanocitta cristata', 'USA'], ['KJ786936', 'Cyanocitta cristata', 'USA'], ['KC333377', 'Cyanocitta cristata', 'USA'], ['KC333380', 'Cyanocitta cristata', 'USA'], ['KJ501437', 'Corvus brachyrhynchos', 'USA']]

Ecotype 15: [['KX547227', 'Culex', 'USA'], ['KX547376', 'Culex', 'USA'], ['KX547358', 'Culex', 'USA'], ['KY216152', 'Culex pipiens', 'USA'], ['KX547483', 'Culex', 'USA'], ['KX547473', 'Culex', 'USA'], ['KJ501530', 'Branta canadensis', 'USA'], ['KY216151', 'Culex pipiens', 'USA'], ['KY229069', 'Culex pipiens', 'USA'], ['KX547604', 'Culex', 'USA'], ['KX547524', 'Culex', 'USA'], ['KM012179', 'Homo sapiens', 'USA'], ['KY216155', 'Culex pipiens', 'USA'], ['JQ700439', 'Homo sapiens', 'USA'], ['KX547391', 'Culex', 'USA'], ['KT020853', 'Homo sapiens', 'USA'], ['KX547184', 'Culex', 'USA'], ['KX547456', 'Culex', 'USA'], ['KM012171', 'Homo sapiens', 'USA'], ['KJ501226', 'Culex tarsalis', 'USA'], ['KX547221', 'Culex', 'USA'], ['KX547334', 'Culex', 'USA'], ['KM012178', 'Homo sapiens', 'USA'], ['KC736500', 'Culex quinquefasciatus', 'USA'], ['KM012176', 'Homo sapiens', 'USA'], ['KJ501220', 'Culex tarsalis', 'USA'], ['KJ501225', 'Culex pipiens', 'USA'], ['KC736498', 'Culex quinquefasciatus', 'USA'], ['KC736497', 'Culex quinquefasciatus', 'USA'], ['KC736496', 'Culex quinquefasciatus', 'USA'], ['KC736494', 'Culex quinquefasciatus', 'USA'], ['KC736486', 'Culex quinquefasciatus', 'USA'], ['KM012180', 'Homo sapiens', 'USA'], ['KC711059', 'Culex restuans', 'USA'], ['MG004536', 'Culex quinquefasciatus', 'USA'], ['KC736492', 'Culex quinquefasciatus', 'USA'], ['KC736495', 'Culex quinquefasciatus', 'USA'], ['MG004541', 'Culex quinquefasciatus', 'USA'], ['MG004535', 'Culex tarsalis', 'USA'], ['MG004528', 'Culex tarsalis', 'USA'], ['MG004531', 'Culex tarsalis', 'USA'], ['MG004530', 'Culex tarsalis', 'USA'], ['MG004534', 'Culex quinquefasciatus', 'USA'], ['MG004532', 'Culex quinquefasciatus', 'USA'], ['MG004538', 'Culex quinquefasciatus', 'USA'], ['KC736493', 'Culex quinquefasciatus', 'USA'], ['KC736490', 'Culex quinquefasciatus', 'USA'], ['KM012183', 'Homo sapiens', 'USA'], ['KM012177', 'Homo sapiens', 'USA'], ['KM012187', 'Homo sapiens', 'USA'], ['KJ501432', 'Corvus brachyrhynchos', 'USA'], ['KX547323', 'Culex', 'USA'], ['KY229072', 'Culex pipiens', 'USA'], ['KX547330', 'Culex', 'USA'], ['KM012188', 'Homo sapiens', 'USA'], ['KX547485', 'Culex', 'USA'], ['KX547196', 'Culex', 'USA'], ['KM012184', 'Homo sapiens', 'USA'], ['KX547373', 'Corvus brachyrhynchos', 'USA'], ['KM012186', 'Homo sapiens', 'USA'], ['KX547557', 'Cyanocitta cristata', 'USA'], ['KX547425', 'Cyanocitta cristata', 'USA'], ['KX547372', 'Corvus brachyrhynchos', 'USA'], ['KX547407', 'Cyanocitta cristata', 'USA'], ['JF920747', 'Culex salinarius', 'USA']]

Ecotype 16: [['KX547475', 'Corvus brachyrhynchos', 'USA'], ['HQ671727', 'Corvus brachyrhynchos', 'USA'], ['HM488196', 'Corvus brachyrhynchos', 'USA'], ['JN183892', 'Culex pipiens', 'USA']]

Ecotype 17: [['KX547424', 'Corvus brachyrhynchos', 'USA'], ['KX547257', 'Culex', 'USA'], ['KX547335', 'Culex', 'USA'], ['KJ501208', 'Culex tarsalis', 'USA'], ['HM488203', 'Corvus brachyrhynchos', 'USA']]

Ecotype 18: [['JF415916', 'Mimus polyglottos', 'USA']]

Ecotype 19: [['KX547284', 'Culex', 'USA'], ['JF415928', 'Culex quinquefasciatus', 'USA'], ['JF415925', 'Aedes albopictus', 'USA'], ['JF415923', 'Culex quinquefasciatus', 'USA'], ['JF415926', 'Culex quinquefasciatus', 'USA'], ['JF415927', 'Aedes albopictus', 'USA'], ['JF415922', 'Culex quinquefasciatus', 'USA'], ['HM488205', 'Corvus brachyrhynchos', 'USA'], ['KJ501219', 'Culex pipiens', 'USA'], ['KJ501207', 'Culex tarsalis', 'USA']]

Ecotype 20: [['HM488243', 'Corvus brachyrhynchos', 'USA'], ['KX547309', 'Corvus brachyrhynchos', 'USA'], ['JF488097', 'Corvus brachyrhynchos', 'USA'], ['HM488199', 'Corvus brachyrhynchos', 'USA'], ['KX547297', 'Corvus brachyrhynchos', 'USA'], ['KC333383', 'Cyanocitta cristata', 'USA'], ['KX547517', 'Corvus brachyrhynchos', 'USA'], ['KX547518', 'Corvus brachyrhynchos', 'USA'], ['KX547494', 'Corvus brachyrhynchos', 'USA'], ['KJ145828', 'Corvus brachyrhynchos', 'USA'], ['HM488165', 'Culex pipiens', 'USA'], ['KC711057', 'Culex quinquefasciatus', 'USA'], ['KJ501360', 'Cyanocitta cristata', 'USA'], ['JF920730', 'Culex pipiens', 'USA']]

Ecotype 21: [['KX547273', 'Culex', 'USA'], ['KX547352', 'Culex', 'USA'], ['KX547451', 'Corvus brachyrhynchos', 'USA'], ['KJ501354', 'Corvus brachyrhynchos', 'USA'], ['KX547282', 'Cyanocitta cristata', 'USA'], ['KX547264', 'Corvus brachyrhynchos', 'USA'], ['KJ501522', 'Corvus brachyrhynchos', 'USA'], ['KX547552', 'Cyanocitta cristata', 'USA'], ['KX547585', 'Corvus brachyrhynchos', 'USA'], ['DQ164203', 'Corvidae', 'USA']]

Ecotype 22: [['KJ145829', 'Corvus brachyrhynchos', 'USA'], ['KX547539', 'Corvus brachyrhynchos', 'USA'], ['HM488201', 'Corvus brachyrhynchos', 'USA'], ['JX015518', 'Culex quinquefasciatus', 'Mexico'], ['JX015520', 'Culex quinquefasciatus', 'Mexico'], ['JX015517', 'Culex tarsalis', 'USA']]

Ecotype 23: [['KX547173', 'Corvus brachyrhynchos', 'USA'], ['HM756675', 'Corvus brachyrhynchos', 'USA'], ['KX547347', 'Corvus brachyrhynchos', 'USA'], ['DQ431711', 'Homo sapiens', 'USA'], ['DQ080051', 'Culex tarsalis', 'USA'], ['GQ507479', 'Homo sapiens', 'USA'], ['DQ080052', 'Culex tarsalis', 'USA'], ['KX547510', 'Culex', 'USA'], ['MG004537', 'Culex quinquefasciatus', 'USA'], ['KJ501108', 'Culex tarsalis', 'USA'], ['KJ501114', 'Culex tarsalis', 'USA'], ['KJ501113', 'Culex tarsalis', 'USA'], ['KJ501112', 'Culex tarsalis', 'USA'], ['GQ507471', 'Homo sapiens', 'USA'], ['MG004533', 'Culex quinquefasciatus', 'USA'], ['MG004540', 'Culex quinquefasciatus', 'USA'], ['JX015523', 'Culex tarsalis', 'USA'], ['KM012172', 'Homo sapiens', 'USA'], ['GQ507470', 'Homo sapiens', 'USA'], ['JX015516', 'Culex tarsalis', 'USA'], ['JX015515', 'Culex tarsalis', 'USA'], ['DQ431702', 'Homo sapiens', 'USA'], ['GQ507468', 'Homo sapiens', 'USA'], ['DQ431712', 'Homo sapiens', 'USA'], ['GQ507482', 'Homo sapiens', 'USA'], ['JF703163', 'Culex tarsalis', 'USA'], ['KJ501106', 'Culex tarsalis', 'USA'], ['KJ501109', 'Culex tarsalis', 'USA'], ['KF704147', 'Culex quinquefasciatus', 'USA'], ['KJ501117', 'Culex tarsalis', 'USA'], ['KJ501120', 'Culex tarsalis', 'USA'], ['KJ501118', 'Culex tarsalis', 'USA'], ['KJ501122', 'Culex tarsalis', 'USA'], ['KJ501121', 'Culex tarsalis', 'USA'], ['KJ501124', 'Culex tarsalis', 'USA'], ['DQ164201', 'Homo sapiens', 'USA'], ['KJ501128', 'Culex quinquefasciatus', 'USA'], ['GQ507469', 'Homo sapiens', 'USA'], ['KJ501102', 'Culex tarsalis', 'USA'], ['KJ501101', 'Culex pipiens', 'USA'], ['KJ501099', 'Culex tarsalis', 'USA'], ['KJ501126', 'Culicidae', 'USA'], ['GQ507484', 'Homo sapiens', 'USA'], ['KJ501111', 'Culex tarsalis', 'USA'], ['KJ501110', 'Culex tarsalis', 'USA'], ['KJ501168', 'Culex quinquefasciatus', 'USA'], ['GQ379159', 'Sciuridae', 'USA'], ['GQ507480', 'Homo sapiens', 'USA'], ['DQ080053', 'Culex tarsalis', 'USA'], ['JF415919', 'Aedes albopictus', 'USA'], ['DQ431704', 'Homo sapiens', 'USA'], ['KJ501478', 'Cyanocitta cristata', 'USA'], ['HM488191', 'Corvus brachyrhynchos', 'USA'], ['DQ431707', 'Homo sapiens', 'USA'], ['DQ431706', 'Homo sapiens', 'USA'], ['DQ080069', 'Equus caballus', 'Mexico'], ['DQ080063', 'Columbidae', 'Mexico'], ['DQ080065', 'Quiscalus', 'Mexico'], ['DQ080068', 'Ardeidae', 'Mexico'], ['DQ080066', 'Phalacrocoracidae', 'Mexico'], ['DQ080067', 'Butorides virescens', 'Mexico'], ['DQ080064', 'Fulica', 'Mexico'], ['DQ080070', 'Quiscalus', 'Mexico'], ['KJ501462', 'Buteo jamaicensis', 'USA'], ['HM488239', 'Corvus brachyrhynchos', 'USA'], ['HM488198', 'Culex pipiens', 'USA'], ['JN183886', 'Cyanocitta cristata', 'USA'], ['KF704153', 'Culex quinquefasciatus', 'USA'], ['JQ700438', 'Homo sapiens', 'USA'], ['KJ501170', 'Culex tarsalis', 'USA'], ['KX547462', 'Corvus brachyrhynchos', 'USA'], ['JF920751', 'Culex restuans', 'USA'], ['JF415921', 'Cyanocitta cristata', 'USA'], ['KX547197', 'Corvus brachyrhynchos', 'USA'], ['KX547187', 'Corvus brachyrhynchos', 'USA'], ['KX547318', 'Cyanocitta cristata', 'USA'], ['JF415917', 'Cyanocitta cristata', 'USA'], ['KJ501295', 'Cyanocitta cristata', 'USA'], ['JF415914', 'Culex quinquefasciatus', 'USA'], ['KX547361', 'Culex', 'USA'], ['KJ501222', 'Culex', 'USA'], ['KC333375', 'Passer domesticus', 'USA']]

Ecotype 24: [['JF730042', 'Corvus brachyrhynchos', 'USA'], ['JF972636', 'Culiseta melanura', 'USA'], ['KX547530', 'Passer domesticus', 'USA'], ['KX547370', 'Culex', 'USA'], ['KX547362', 'Culex', 'USA'], ['KX547219', 'Culex', 'USA'], ['KX547208', 'Culex', 'USA'], ['HM488242', 'Poecile atricapilla', 'USA'], ['KX547191', 'Culex', 'USA'], ['HM488177', 'Corvus brachyrhynchos', 'USA'], ['DQ164205', 'Homo sapiens', 'USA'], ['DQ164198', 'Homo sapiens', 'USA'], ['KX547588', 'Turdus migratorius', 'USA'], ['KX547505', 'Culex', 'USA'], ['KJ501331', 'Buteo jamaicensis', 'USA'], ['HM488254', 'Culex', 'USA'], ['KX547289', 'Corvus brachyrhynchos', 'USA'], ['KX547549', 'Corvus brachyrhynchos', 'USA'], ['HM488251', 'Corvus brachyrhynchos', 'USA'], ['HQ671728', 'Corvus brachyrhynchos', 'USA'], ['KJ501266', 'Corvus brachyrhynchos', 'USA'], ['DQ164191', 'Corvus brachyrhynchos', 'USA'], ['HM756648', 'Ochlerotatus trivittatus', 'USA']]

Ecotype 25: [['KJ501466', 'Bubo virginianus', 'USA'], ['KX547459', 'Corvus brachyrhynchos', 'USA']]

Ecotype 26: [['KX547292', 'Culex', 'USA'], ['JF415920', 'Cyanocitta cristata', 'USA'], ['KX547300', 'Corvus brachyrhynchos', 'USA']]

Ecotype 27: [['DQ164200', 'Homo sapiens', 'USA'], ['KX547189', 'Culex', 'USA'], ['JF899528', 'Corvus brachyrhynchos', 'USA']]

Ecotype 28: [['KX547527', 'Equus caballus', 'USA'], ['KX547402', 'Culex', 'USA'], ['KX547204', 'Corvus brachyrhynchos', 'USA'], ['JF920744', 'Culex pipiens', 'USA'], ['KX547481', 'Cyanocitta cristata', 'USA'], ['HM756668', 'Corvus brachyrhynchos', 'USA'], ['KX547526', 'Culex', 'USA'], ['KX547531', 'Culex', 'USA'], ['HM756652', 'Aedes cinereus', 'USA'], ['KX547377', 'Culex', 'USA'], ['KX547621', 'Cyanocitta cristata', 'USA']]

Ecotype 29: [['KX547525', 'Corvus brachyrhynchos', 'USA']]

Ecotype 30: [['KX547503', 'Culex', 'USA'], ['HQ671742', 'Cyanocitta cristata', 'USA'], ['KJ501336', 'Cyanocitta cristata', 'USA'], ['KX547218', 'Culex', 'USA'], ['KJ501464', 'Cyanocitta cristata', 'USA'], ['HM756667', 'Corvus brachyrhynchos', 'USA'], ['HM488253', 'Culex', 'USA'], ['KX547211', 'Corvus brachyrhynchos', 'USA'], ['KX547500', 'Corvus brachyrhynchos', 'USA'], ['KX547535', 'Cyanocitta cristata', 'USA'], ['KX547290', 'Corvus brachyrhynchos', 'USA'], ['KX547442', 'Culex', 'USA'], ['JF920740', 'Culex pipiens', 'USA'], ['JF920748', 'Culex pipiens', 'USA'], ['JF920742', 'Culex pipiens', 'USA'], ['JF920755', 'Culex pipiens', 'USA'], ['KX547543', 'Culex', 'USA'], ['KX547458', 'Cyanocitta cristata', 'USA'], ['KM012173', 'Homo sapiens', 'USA'], ['KX547419', 'Corvus brachyrhynchos', 'USA'], ['HM488200', 'Corvus brachyrhynchos', 'USA']]

Ecotype 31: [['KX547427', 'Cyanocitta cristata', 'USA'], ['KX547610', 'Culex', 'USA'], ['KX547477', 'Culex', 'USA'], ['KX547357', 'Culex', 'USA'], ['DQ164195', 'Culex', 'USA'], ['KJ501366', 'Corvus brachyrhynchos', 'USA'], ['HM756673', 'Corvus brachyrhynchos', 'USA']]

Ecotype 32: [['HQ671722', 'Corvus brachyrhynchos', 'USA']]

Ecotype 33: [['KX547262', 'Corvus brachyrhynchos', 'USA']]

Ecotype 34: [['KJ501292', 'Corvus brachyrhynchos', 'USA'], ['KJ501361', 'Cyanocitta cristata', 'USA'], ['JN819320', 'Aves', 'USA'], ['HM488211', 'Culex restuans', 'USA'], ['HM488138', 'Culex restuans', 'USA'], ['KX547288', 'Corvus brachyrhynchos', 'USA'], ['HM488235', 'Culiseta melanura', 'USA'], ['HQ671699', 'Ochlerotatus sollicitans', 'USA'], ['HM488137', 'Culex pipiens', 'USA'], ['HQ671698', 'Culex restuans', 'USA']]

Ecotype 35: [['KX547310', 'Culex', 'USA'], ['HM756654', 'Culex salinarius', 'USA'], ['KJ501498', 'Passer domesticus', 'USA'], ['JF488086', 'Culex pipiens', 'USA'], ['JF488092', 'Culex restuans', 'USA'], ['KX547267', 'Cyanocitta cristata', 'USA'], ['JF488089', 'Culex restuans', 'USA'], ['KX547396', 'Culex', 'USA'], ['KX547366', 'Carpodacus mexicanus', 'USA'], ['KX547332', 'Corvus brachyrhynchos', 'USA'], ['KX547217', 'Colaptes auratus', 'USA'], ['KX547307', 'Cyanocitta cristata', 'USA'], ['KX547303', 'Culex', 'USA'], ['JF920745', 'Culex pipiens', 'USA'], ['JF488087', 'Culex pipiens', 'USA'], ['KX547590', 'Carpodacus mexicanus', 'USA'], ['KX547486', 'Culex', 'USA'], ['KX547392', 'Culex', 'USA'], ['KX547496', 'Corvus brachyrhynchos', 'USA'], ['KX547553', 'Cyanocitta cristata', 'USA'], ['JF920760', 'Culex pipiens', 'USA'], ['GQ507481', 'Homo sapiens', 'USA'], ['KX547521', 'Corvus brachyrhynchos', 'USA'], ['JF488090', 'Culex pipiens', 'USA'], ['KX547597', 'Cyanocitta cristata', 'USA'], ['KX547256', 'Culex', 'USA'], ['KX547308', 'Cyanocitta cristata', 'USA'], ['HQ671700', 'Culiseta melanura', 'USA'], ['KX547455', 'Culex', 'USA'], ['KJ501273', 'Cyanocitta cristata', 'USA'], ['JF920738', 'Culex pipiens', 'USA']]

Ecotype 36: [['KX547563', 'Culex', 'USA'], ['KX547238', 'Corvus brachyrhynchos', 'USA'], ['KX547341', 'Corvus brachyrhynchos', 'USA'], ['KX547314', 'Culex', 'USA'], ['KX547245', 'Culex', 'USA'], ['KX547331', 'Cyanocitta cristata', 'USA'], ['KX547513', 'Culex', 'USA']]

Ecotype 37: [['KJ501456', 'Corvus brachyrhynchos', 'USA'], ['KJ501457', 'Corvus brachyrhynchos', 'USA'], ['KX547363', 'Culex', 'USA'], ['KX547312', 'Corvus brachyrhynchos', 'USA'], ['JN183885', 'Cyanocitta cristata', 'USA'], ['KX547606', 'Culex', 'USA']]

Ecotype 38: [['KJ501345', 'Corvus brachyrhynchos', 'USA'], ['HM488230', 'Culex salinarius', 'USA'], ['KJ501257', 'Corvus brachyrhynchos', 'USA'], ['KX547446', 'Corvus brachyrhynchos', 'USA'], ['KX547599', 'Culex', 'USA'], ['JF920735', 'Culex pipiens', 'USA'], ['HQ671726', 'Corvus brachyrhynchos', 'USA'], ['KX547360', 'Corvus brachyrhynchos', 'USA'], ['KX547349', 'Culex', 'USA'], ['JF920733', 'Culex pipiens', 'USA'], ['KX547561', 'Corvus brachyrhynchos', 'USA'], ['KX547545', 'Corvus brachyrhynchos', 'USA'], ['KX547215', 'Culex', 'USA'], ['JF415929', 'Cyanocitta cristata', 'USA'], ['HM488197', 'Corvus brachyrhynchos', 'USA'], ['KX547436', 'Culex', 'USA'], ['HM488241', 'Corvus brachyrhynchos', 'USA'], ['KX547580', 'Corvus brachyrhynchos', 'USA'], ['HM488184', 'Cyanocitta cristata', 'USA'], ['HM488210', 'Culiseta melanura', 'USA'], ['KX547592', 'Culex', 'USA'], ['GU828004', 'Cyanocitta cristata', 'USA'], ['GU828001', 'Culex quinquefasciatus', 'USA'], ['GU828003', 'Zenaida macroura', 'USA'], ['GU828002', 'Culex quinquefasciatus', 'USA'], ['AY712948', 'Culex quinquefasciatus', 'USA'], ['KJ501321', 'Corvus brachyrhynchos', 'USA'], ['KJ501384', 'Cyanocitta cristata', 'USA'], ['KX547605', 'Corvus brachyrhynchos', 'USA'], ['HM488193', 'Corvus brachyrhynchos', 'USA'], ['KJ501309', 'Cyanocitta cristata', 'USA'], ['KX547615', 'Corvus brachyrhynchos', 'USA'], ['HM756649', 'Culex pipiens', 'USA'], ['HM488160', 'Culex pipiens', 'USA'], ['KJ501368', 'Corvus brachyrhynchos', 'USA'], ['HQ705660', 'Corvus brachyrhynchos', 'USA'], ['HM488173', 'Culex restuans', 'USA'], ['HM488217', 'Culex salinarius', 'USA'], ['HM756650', 'Culex salinarius', 'USA']]

Ecotype 39: [['JN183891', 'Cyanocitta cristata', 'USA'], ['KJ501458', 'Cyanocitta cristata', 'USA'], ['KX547336', 'Corvus brachyrhynchos', 'USA'], ['DQ431693', 'Homo sapiens', 'USA'], ['KX547188', 'Culex', 'USA'], ['KX547593', 'Cyanocitta cristata', 'USA'], ['KJ501387', 'Cyanocitta cristata', 'USA'], ['KX547182', 'Culex', 'USA'], ['KX547226', 'Culex', 'USA'], ['KX547275', 'Cyanocitta cristata', 'USA'], ['KJ501284', 'Cyanocitta cristata', 'USA'], ['HM488212', 'Culex salinarius', 'USA'], ['HM488176', 'Culex salinarius', 'USA'], ['KX547231', 'Culex', 'USA'], ['KX547260', 'Corvus brachyrhynchos', 'USA'], ['KJ501511', 'Cyanocitta cristata', 'USA'], ['HM488179', 'Corvus brachyrhynchos', 'USA'], ['HM756651', 'Ochlerotatus trivittatus', 'USA'], ['KJ501451', 'Corvus brachyrhynchos', 'USA'], ['KX547345', 'Corvus brachyrhynchos', 'USA']]

Ecotype 40: [['KJ501503', 'Turdus migratorius', 'USA'], ['KJ501504', 'Branta canadensis', 'USA'], ['KX547506', 'Corvus brachyrhynchos', 'USA'], ['KJ501271', 'Bubo virginianus', 'USA'], ['HM488144', 'Culex restuans', 'USA'], ['HM488146', 'Psorophora ferox', 'USA'], ['HM488145', 'Aedes vexans', 'USA'], ['JF920749', 'Culex pipiens', 'USA'], ['JF920739', 'Culex pipiens', 'USA'], ['JF920729', 'Culex pipiens', 'USA'], ['JF920737', 'Culex pipiens', 'USA'], ['JF920732', 'Culex pipiens', 'USA'], ['JF920736', 'Culex pipiens', 'USA'], ['KX547225', 'Culex', 'USA'], ['KX547294', 'Culex', 'USA'], ['HM488172', 'Ochlerotatus sticticus', 'USA'], ['HM488223', 'Culiseta melanura', 'USA'], ['HM488224', 'Culiseta melanura', 'USA'], ['KX547421', 'Corvus brachyrhynchos', 'USA'], ['KX547603', 'Corvus brachyrhynchos', 'USA'], ['KJ501411', 'Passer domesticus', 'USA'], ['KJ501412', 'Passer domesticus', 'USA'], ['HM756676', 'Cyanocitta cristata', 'USA'], ['DQ431699', 'Homo sapiens', 'USA'], ['KX547241', 'Culex', 'USA'], ['KJ501312', 'Corvus brachyrhynchos', 'USA'], ['KX547512', 'Corvus brachyrhynchos', 'USA'], ['KX547476', 'Cyanocitta cristata', 'USA'], ['KX547175', 'Culex', 'USA'], ['KX547447', 'Corvus brachyrhynchos', 'USA'], ['KX547453', 'Turdus migratorius', 'USA'], ['HM488204', 'Corvus brachyrhynchos', 'USA'], ['KX547463', 'Culex', 'USA'], ['KX547541', 'Culex', 'USA'], ['KX547164', 'Culex', 'USA'], ['JF415915', 'Quiscalus quiscula', 'USA'], ['JF415930', 'Culex quinquefasciatus', 'USA'], ['KJ501380', 'Bubo virginianus', 'USA'], ['HM488159', 'Culex salinarius', 'USA'], ['KX547259', 'Cyanocitta cristata', 'USA'], ['KJ786935', 'Mimus polyglottos', 'USA'], ['KJ501303', 'Cyanocitta cristata', 'USA'], ['HM488188', 'Corvus brachyrhynchos', 'USA'], ['HM488178', 'Corvus brachyrhynchos', 'USA'], ['DQ164193', 'Corvus brachyrhynchos', 'USA'], ['KX547305', 'Corvus brachyrhynchos', 'USA'], ['JF730043', 'Culex pipiens', 'USA'], ['KX547618', 'Corvus brachyrhynchos', 'USA'], ['HM756669', 'Corvus brachyrhynchos', 'USA'], ['KX547448', 'Corvus brachyrhynchos', 'USA'], ['DQ431696', 'Homo sapiens', 'USA'], ['KX547422', 'Cyanocitta cristata', 'USA'], ['HQ671720', 'Corvus brachyrhynchos', 'USA'], ['KX547302', 'Corvus brachyrhynchos', 'USA'], ['KX547346', 'Corvus brachyrhynchos', 'USA'], ['KJ501376', 'Bubo virginianus', 'USA'], ['KX547213', 'Cyanocitta cristata', 'USA'], ['HM488233', 'Aedes vexans', 'USA'], ['HM488234', 'Culex salinarius', 'USA'], ['HM488240', 'Cyanocitta cristata', 'USA'], ['HQ671725', 'Corvus brachyrhynchos', 'USA'], ['KX547508', 'Corvus brachyrhynchos', 'USA'], ['KJ501296', 'Pica pica', 'USA'], ['JX015521', 'Culex tarsalis', 'USA'], ['JX015519', 'Culex quinquefasciatus', 'USA'], ['JX015522', 'Culex tarsalis', 'USA'], ['KJ501217', 'Culex tarsalis', 'USA'], ['KX547276', 'Culex', 'USA'], ['KJ501475', 'Cyanocitta cristata', 'USA'], ['KJ501315', 'Corvus brachyrhynchos', 'USA'], ['JF415924', 'Cyanocitta cristata', 'USA'], ['KJ501214', 'Culex tarsalis', 'USA'], ['KJ501254', 'Branta canadensis', 'USA'], ['KX547472', 'Cyanocitta cristata', 'USA'], ['KX547608', 'Culex', 'USA'], ['KT862844', 'Corvus brachyrhynchos', 'USA'], ['KT862843', 'Cyanocitta cristata', 'USA'], ['KC333374', 'Passer domesticus', 'USA'], ['KX547168', 'Carpodacus mexicanus', 'USA'], ['HM488237', 'Corvus brachyrhynchos', 'USA'], ['KX547247', 'Culex', 'USA']]

Ecotype 41: [['JN819311', 'Culicidae', 'USA'], ['HQ671703', 'Culiseta melanura', 'USA'], ['JF920728', 'Culex pipiens', 'USA'], ['KX547287', 'Passer domesticus', 'USA'], ['KJ501527', 'Corvus brachyrhynchos', 'USA'], ['KX547207', 'Corvus brachyrhynchos', 'USA'], ['KX547311', 'Sciurus carolinensis', 'USA'], ['KX547609', 'Sciurus carolinensis', 'USA'], ['HM488220', 'Culex salinarius', 'USA'], ['KJ501447', 'Corvus brachyrhynchos', 'USA'], ['KJ501396', 'Corvus brachyrhynchos', 'USA'], ['HM488164', 'Culex pipiens', 'USA'], ['HM488161', 'Culex pipiens', 'USA'], ['HM488163', 'Culex pipiens', 'USA'], ['JF920750', 'Culex pipiens', 'USA'], ['KX547401', 'Culex', 'USA'], ['KX547265', 'Cyanocitta cristata', 'USA'], ['KX547440', 'Corvus brachyrhynchos', 'USA'], ['KX547365', 'Cyanocitta cristata', 'USA'], ['KX547611', 'Corvus brachyrhynchos', 'USA'], ['KX547568', 'Corvus brachyrhynchos', 'USA'], ['KX547498', 'Corvus brachyrhynchos', 'USA'], ['HM488228', 'Culex salinarius', 'USA'], ['HM488227', 'Culex restuans', 'USA'], ['KJ501351', 'Corvus brachyrhynchos', 'USA'], ['HQ705669', 'Cyanocitta cristata', 'USA']]

Ecotype 42: [['DQ431695', 'Homo sapiens', 'USA'], ['HM488155', 'Culex restuans', 'USA'], ['KX547468', 'Culex', 'USA'], ['GU827999', 'Cyanocitta cristata', 'USA'], ['KX547388', 'Culex', 'USA'], ['GU828000', 'Cyanocitta cristata', 'USA'], ['DQ164199', 'Homo sapiens', 'USA'], ['HQ671721', 'Corvus brachyrhynchos', 'USA']]

Ecotype 43: [['KJ501265', 'Cyanocitta cristata', 'USA'], ['KJ501373', 'Cyanocitta cristata', 'USA'], ['KJ501369', 'Passer domesticus', 'USA']]

Ecotype 44: [['KX547246', 'Corvus brachyrhynchos', 'USA'], ['KJ501514', 'Corvus brachyrhynchos', 'USA'], ['DQ164196', 'Homo sapiens', 'USA'], ['DQ164197', 'Homo sapiens', 'USA']]

Ecotype 45: [['HM488123', 'Culex pipiens', 'USA'], ['HM488124', 'Mesocricetus auratus', 'USA'], ['HM488122', 'Culex pipiens', 'USA'], ['HM488121', 'Culex pipiens', 'USA'], ['HM488215', 'Culiseta melanura', 'USA'], ['KX547620', 'Corvus brachyrhynchos', 'USA'], ['KX547166', 'Equus caballus', 'USA'], ['JN367277', 'Corvus brachyrhynchos', 'USA'], ['HM756665', 'Corvus brachyrhynchos', 'USA'], ['KX547586', 'Culex', 'USA']]

Ecotype 46: [['KJ501339', 'Cyanocitta cristata', 'USA'], ['KJ501280', 'Corvus brachyrhynchos', 'USA'], ['HM488226', 'Culex pipiens', 'USA'], ['HM488225', 'Aedes cinereus', 'USA'], ['HM488229', 'Psorophora ferox', 'USA'], ['HM488185', 'Cyanocitta cristata', 'USA'], ['HM488143', 'Aedes cinereus', 'USA'], ['KJ501287', 'Cyanocitta cristata', 'USA'], ['KJ501337', 'Corvus brachyrhynchos', 'USA'], ['KJ501472', 'Corvus brachyrhynchos', 'USA']]

Ecotype 47: [['HM488187', 'Corvus brachyrhynchos', 'USA'], ['KJ501251', 'Corvus brachyrhynchos', 'USA'], ['KJ501300', 'Cyanocitta cristata', 'USA'], ['DQ164204', 'Buteo jamaicensis', 'USA'], ['KJ501329', 'Branta canadensis', 'USA'], ['KJ501388', 'Cyanocitta cristata', 'USA'], ['KJ501502', 'Corvus brachyrhynchos', 'USA'], ['KJ501347', 'Corvus brachyrhynchos', 'USA'], ['DQ080055', 'Culex tarsalis', 'USA'], ['JF703164', 'Culex tarsalis', 'USA'], ['KJ501105', 'Culex tarsalis', 'USA'], ['JF703162', 'Culex tarsalis', 'USA'], ['DQ080056', 'Culex tarsalis', 'USA'], ['JF703161', 'Culex tarsalis', 'USA'], ['DQ431700', 'Homo sapiens', 'USA'], ['KJ501123', 'Culex tarsalis', 'USA'], ['DQ431708', 'Homo sapiens', 'USA'], ['KJ501125', 'Culex tarsalis', 'USA'], ['KJ501104', 'Culex tarsalis', 'USA'], ['KJ501103', 'Culex tarsalis', 'USA'], ['KJ501107', 'Culex tarsalis', 'USA'], ['KJ501186', 'Culex tarsalis', 'USA'], ['DQ080060', 'Corvus', 'Mexico'], ['GQ507473', 'Homo sapiens', 'USA'], ['KJ501129', 'Culex quinquefasciatus', 'USA'], ['KJ501116', 'Culex quinquefasciatus', 'USA'], ['GQ379157', 'Corvus', 'USA'], ['KJ501143', 'Culex quinquefasciatus', 'USA'], ['KJ501127', 'Culicidae', 'USA'], ['KJ501135', 'Culex quinquefasciatus', 'USA'], ['KJ501136', 'Culex quinquefasciatus', 'USA'], ['KJ501138', 'Culex quinquefasciatus', 'USA'], ['KJ501140', 'Culex quinquefasciatus', 'USA'], ['KJ501141', 'Culex quinquefasciatus', 'USA'], ['KJ501201', 'Culex quinquefasciatus', 'USA'], ['KJ501137', 'Culex quinquefasciatus', 'USA'], ['KJ501142', 'Culicidae', 'USA'], ['KJ501130', 'Culex quinquefasciatus', 'USA'], ['KJ501150', 'Culex quinquefasciatus', 'USA'], ['KJ501148', 'Culicidae', 'USA'], ['KJ501149', 'Culex quinquefasciatus', 'USA'], ['KJ501134', 'Culex quinquefasciatus', 'USA'], ['KJ501133', 'Culex quinquefasciatus', 'USA'], ['GQ379158', 'Culex tarsalis', 'USA'], ['KJ501147', 'Culex quinquefasciatus', 'USA'], ['KJ501144', 'Culex quinquefasciatus', 'USA'], ['KJ501139', 'Culex quinquefasciatus', 'USA'], ['KJ501146', 'Culex quinquefasciatus', 'USA'], ['GQ507474', 'Homo sapiens', 'USA'], ['GQ507477', 'Homo sapiens', 'USA'], ['GQ507478', 'Homo sapiens', 'USA'], ['GQ507483', 'Homo sapiens', 'USA'], ['GQ507476', 'Homo sapiens', 'USA'], ['KJ501132', 'Culex quinquefasciatus', 'USA'], ['DQ080054', 'Culex quinquefasciatus', 'USA'], ['DQ431709', 'Homo sapiens', 'USA'], ['KJ501119', 'Culex tarsalis', 'USA'], ['GQ507475', 'Homo sapiens', 'USA'], ['KJ501115', 'Culex tarsalis', 'USA'], ['KJ501131', 'Culex quinquefasciatus', 'USA'], ['DQ431710', 'Homo sapiens', 'USA'], ['KJ501178', 'Culex tarsalis', 'USA'], ['JQ700441', 'Homo sapiens', 'USA'], ['KM012175', 'Homo sapiens', 'USA'], ['KJ501100', 'Culex tarsalis', 'USA'], ['KJ501098', 'Culex pipiens', 'USA'], ['KJ501096', 'Culex tarsalis', 'USA'], ['DQ080059', 'Corvidae', 'USA'], ['KJ501206', 'Culex tarsalis', 'USA'], ['KJ501195', 'Culex tarsalis', 'USA'], ['DQ080058', 'Corvus', 'USA'], ['DQ080057', 'Corvus', 'USA'], ['DQ176637', 'Quiscalus quiscula', 'USA'], ['GU827998', 'Cyanocitta cristata', 'USA'], ['AY712945', 'Zenaida macroura', 'USA'], ['AY712946', 'Cyanocitta cristata', 'USA'], ['KJ501501', 'Corvus brachyrhynchos', 'USA'], ['KJ501352', 'Corvus brachyrhynchos', 'USA'], ['KJ501356', 'Cyanocitta cristata', 'USA'], ['HM488208', 'Culex salinarius', 'USA'], ['KJ501311', 'Cyanocitta cristata', 'USA'], ['KJ501481', 'Corvus brachyrhynchos', 'USA'], ['HQ671704', 'Culex pipiens', 'USA'], ['KJ501301', 'Cyanocitta cristata', 'USA'], ['KJ501463', 'Corvus brachyrhynchos', 'USA']]

Ecotype 48: [['JF920746', 'Culex pipiens', 'USA'], ['KX547578', 'Cyanocitta cristata', 'USA'], ['GQ507472', 'Homo sapiens', 'USA'], ['KX547569', 'Corvus brachyrhynchos', 'USA'], ['KX547183', 'Culex', 'USA'], ['DQ431694', 'Homo sapiens', 'USA'], ['KJ501298', 'Cyanocitta cristata', 'USA'], ['HQ671730', 'Corvus brachyrhynchos', 'USA'], ['KX547426', 'Culex', 'USA'], ['HM488162', 'Culex pipiens', 'USA'], ['HM488189', 'Cyanocitta cristata', 'USA'], ['DQ431705', 'Homo sapiens', 'USA'], ['KJ501353', 'Corvus brachyrhynchos', 'USA'], ['KJ501482', 'Cyanocitta cristata', 'USA'], ['KJ501508', 'Zenaida macroura', 'USA'], ['HM488195', 'Cyanocitta cristata', 'USA'], ['JF920756', 'Culex pipiens', 'USA'], ['KJ501474', 'Bubo virginianus', 'USA'], ['HM488180', 'Corvus brachyrhynchos', 'USA'], ['JN819319', 'Aves', 'USA'], ['KX547546', 'Corvus brachyrhynchos', 'USA'], ['KX547354', 'Corvus brachyrhynchos', 'USA'], ['KJ501449', 'Cyanocitta cristata', 'USA'], ['KX547258', 'Corvus brachyrhynchos', 'USA'], ['KX547523', 'Corvus brachyrhynchos', 'USA'], ['KX547397', 'Corvus brachyrhynchos', 'USA'], ['KX547355', 'Cyanocitta cristata', 'USA'], ['KX547270', 'Corvus brachyrhynchos', 'USA'], ['KX547614', 'Culex', 'USA'], ['KX547571', 'Corvus brachyrhynchos', 'USA'], ['KX547379', 'Culex', 'USA'], ['KJ501487', 'Corvus brachyrhynchos', 'USA']]

Ecotype 49: [['KJ501454', 'Corvus brachyrhynchos', 'USA']]

Ecotype 50: [['KJ501283', 'Corvus brachyrhynchos', 'USA'], ['KX547384', 'Carpodacus mexicanus', 'USA'], ['HM488114', 'Aedes cinereus', 'USA'], ['HM488139', 'Culex salinarius', 'USA'], ['HQ705659', 'Aedes cinereus', 'USA'], ['HM488141', 'Culex pipiens', 'USA'], ['KX547304', 'Corvus brachyrhynchos', 'USA'], ['KX547420', 'Cyanocitta cristata', 'USA'], ['KX547271', 'Cyanocitta cristata', 'USA'], ['KJ501397', 'Cyanocitta cristata', 'USA'], ['JN183887', 'Corvus brachyrhynchos', 'USA'], ['KX547181', 'Cyanocitta cristata', 'USA'], ['KJ501519', 'Corvus brachyrhynchos', 'USA'], ['KJ501333', 'Cyanocitta cristata', 'USA'], ['KJ501490', 'Corvus brachyrhynchos', 'USA'], ['KX547268', 'Culex', 'USA'], ['KX547339', 'Cyanocitta cristata', 'USA'], ['HM488202', 'Cyanocitta cristata', 'USA'], ['HM488207', 'Cyanocitta cristata', 'USA'], ['KX547558', 'Culex', 'USA'], ['KX547504', 'Culex', 'USA'], ['KX547348', 'Culex', 'USA'], ['KX547434', 'Culex', 'USA'], ['KX547452', 'Corvus brachyrhynchos', 'USA'], ['HM488245', 'Corvus brachyrhynchos', 'USA'], ['KJ501281', 'Cyanocitta cristata', 'USA'], ['HM488149', 'Ochlerotatus cantator', 'USA'], ['HM488152', 'Aedes vexans', 'USA'], ['HM488115', 'Culex salinarius', 'USA'], ['HM488118', 'Culex pipiens', 'USA'], ['HM488117', 'Ochlerotatus triseriatus', 'USA'], ['HM488119', 'Culex pipiens', 'USA'], ['HM488151', 'Culex pipiens', 'USA'], ['HM488154', 'Culex pipiens', 'USA'], ['HM488120', 'Culex pipiens', 'USA'], ['HM488150', 'Culex pipiens', 'USA'], ['HM488153', 'Culex pipiens', 'USA'], ['HM756656', 'Culiseta melanura', 'USA'], ['DQ431698', 'Homo sapiens', 'USA'], ['JF488091', 'Culex salinarius', 'USA'], ['KJ501365', 'Corvus brachyrhynchos', 'USA'], ['JN819312', 'Culicidae', 'USA'], ['HM488157', 'Culex pipiens', 'USA'], ['HM488221', 'Culiseta melanura', 'USA'], ['HM488213', 'Culex restuans', 'USA'], ['KJ501272', 'Cyanocitta cristata', 'USA'], ['JN819313', 'Culicidae', 'USA'], ['KJ501460', 'Sciurus carolinensis', 'USA'], ['HM488148', 'Culex pipiens', 'USA'], ['HM488116', 'Culex pipiens', 'USA'], ['HM488156', 'Culex pipiens', 'USA'], ['HM488158', 'Culex pipiens', 'USA']]

Ecotype 51: [['KJ501452', 'Corvus brachyrhynchos', 'USA'], ['KJ501477', 'Corvus brachyrhynchos', 'USA'], ['KJ501328', 'Buteo jamaicensis', 'USA'], ['KX547488', 'Culex', 'USA'], ['DQ164189', 'Corvus brachyrhynchos', 'USA'], ['HM756657', 'Culex pipiens', 'USA'], ['KX547489', 'Corvus brachyrhynchos', 'USA'], ['KX547232', 'Passer domesticus', 'USA'], ['HM488171', 'Culex restuans', 'USA'], ['JF920306', 'Culex pipiens', 'USA'], ['KJ501484', 'Corvus brachyrhynchos', 'USA'], ['KX547529', 'Culex', 'USA'], ['KX547404', 'Corvus brachyrhynchos', 'USA'], ['KX547449', 'Culex', 'USA'], ['HM488190', 'Corvus brachyrhynchos', 'USA'], ['KJ501506', 'Corvus brachyrhynchos', 'USA'], ['KJ501355', 'Cyanocitta cristata', 'USA'], ['KJ501426', 'Corvus brachyrhynchos', 'USA'], ['KJ501510', 'Cyanocitta cristata', 'USA']]

Ecotype 52: [['KJ501319', 'Corvus brachyrhynchos', 'USA'], ['HM488181', 'Corvus brachyrhynchos', 'USA'], ['KX547536', 'Corvus brachyrhynchos', 'USA'], ['KC333379', 'Cyanocitta cristata', 'USA'], ['KC333382', 'Cyanocitta cristata', 'USA'], ['KC333386', 'Cyanocitta cristata', 'USA'], ['JF920757', 'Culex pipiens', 'USA'], ['KX547278', 'Corvus brachyrhynchos', 'USA'], ['JF920743', 'Culex pipiens', 'USA'], ['KX547248', 'Culex', 'USA'], ['KX547327', 'Corvus brachyrhynchos', 'USA'], ['JN183890', 'Culiseta melanura', 'USA'], ['JN819318', 'Aves', 'USA'], ['JN183888', 'Corvus brachyrhynchos', 'USA'], ['KX547277', 'Culex', 'USA'], ['KC333387', 'Passer domesticus', 'USA'], ['KJ501476', 'Pica pica', 'USA'], ['KX547240', 'Culex', 'USA'], ['JF920741', 'Culex pipiens', 'USA'], ['KX547244', 'Corvus brachyrhynchos', 'USA'], ['KJ501364', 'Corvus brachyrhynchos', 'USA'], ['HM488252', 'Corvus brachyrhynchos', 'USA'], ['KX547501', 'Corvus brachyrhynchos', 'USA'], ['JF488093', 'Culex pipiens', 'USA'], ['KX547441', 'Passer domesticus', 'USA'], ['KX547502', 'Culex', 'USA'], ['JF899529', 'Corvus brachyrhynchos', 'USA'], ['KX547193', 'Culex', 'USA'], ['KX547532', 'Cyanocitta cristata', 'USA'], ['KX547487', 'Corvus brachyrhynchos', 'USA'], ['KX547351', 'Culex', 'USA'], ['KX547567', 'Corvus brachyrhynchos', 'USA'], ['KX547212', 'Culex', 'USA'], ['HM756658', 'Culiseta melanura', 'USA']]

Ecotype 53: [['KJ501467', 'Bubo virginianus', 'USA'], ['KX547381', 'Cyanocitta cristata', 'USA'], ['HM488182', 'Corvus brachyrhynchos', 'USA'], ['KJ501370', 'Cyanocitta cristata', 'USA'], ['KX547408', 'Culex', 'USA'], ['HM488194', 'Corvus brachyrhynchos', 'USA'], ['KX547409', 'Corvus brachyrhynchos', 'USA'], ['KX547577', 'Corvus brachyrhynchos', 'USA'], ['KX547296', 'Corvus brachyrhynchos', 'USA'], ['KX547399', 'Culex', 'USA'], ['HM756660', 'Accipiter cooperii', 'USA'], ['KX547582', 'Culex', 'USA'], ['KX547515', 'Corvus brachyrhynchos', 'USA'], ['KX547491', 'Corvus brachyrhynchos', 'USA'], ['KX547306', 'Culex', 'USA'], ['KX547589', 'Cyanocitta cristata', 'USA'], ['KX547406', 'Culex', 'USA'], ['JF920754', 'Culex pipiens', 'USA'], ['JF920759', 'Culex pipiens', 'USA'], ['KY229074', 'Culex pipiens', 'USA'], ['KX547315', 'Culex', 'USA'], ['KJ501533', 'Culicidae', 'USA'], ['KM012181', 'Homo sapiens', 'USA'], ['KX547465', 'Culex', 'USA'], ['KX547190', 'Culex', 'USA'], ['KX547254', 'Culex', 'USA'], ['KX547235', 'Culex', 'USA'], ['KX547612', 'Culex', 'USA'], ['KY229073', 'Culex pipiens', 'USA'], ['KX547200', 'Culex', 'USA'], ['KX547174', 'Culex', 'USA'], ['KX547337', 'Culex', 'USA'], ['KX547333', 'Culex', 'USA'], ['KY216153', 'Culex pipiens', 'USA'], ['KY229070', 'Culex pipiens', 'USA'], ['KX547520', 'Culex', 'USA'], ['KC333376', 'Cyanocitta cristata', 'USA'], ['KJ501216', 'Culex pipiens', 'USA'], ['KC333381', 'Cyanocitta cristata', 'USA'], ['KC711058', 'Culex quinquefasciatus', 'USA'], ['KC736491', 'Culex quinquefasciatus', 'USA'], ['KC736489', 'Culex quinquefasciatus', 'USA'], ['KC736499', 'Culex quinquefasciatus', 'USA'], ['KC736487', 'Culex quinquefasciatus', 'USA'], ['KC736488', 'Culex quinquefasciatus', 'USA'], ['KC736501', 'Culex quinquefasciatus', 'USA'], ['KC736502', 'Culex quinquefasciatus', 'USA'], ['KJ501230', 'Culex tarsalis', 'USA'], ['JF488095', 'Corvus brachyrhynchos', 'USA'], ['KX547555', 'Culex', 'USA'], ['KC333378', 'Cyanocitta cristata', 'USA'], ['KX547186', 'Culex', 'USA'], ['JF488096', 'Corvus brachyrhynchos', 'USA'], ['KJ501377', 'Bubo virginianus', 'USA'], ['KX547403', 'Corvus brachyrhynchos', 'USA']]

Ecotype 54: [['KJ501453', 'Corvus brachyrhynchos', 'USA'], ['KJ501304', 'Corvus brachyrhynchos', 'USA']]

Ecotype 55: [['AY646354', 'Homo sapiens', 'USA'], ['KX547493', 'Corvus brachyrhynchos', 'USA'], ['HM488147', 'Culex salinarius', 'USA'], ['HM488142', 'Ochlerotatus triseriatus', 'USA'], ['KX547454', 'Culex', 'USA'], ['KX547176', 'Culex', 'USA']]

Ecotype 56: [['AF404755', 'Bonasa umbellus', 'USA'], ['KJ501379', 'Cyanocitta cristata', 'USA'], ['KJ501473', 'Corvus brachyrhynchos', 'USA'], ['DQ983578', 'Culex nigripalpus', 'USA'], ['KJ501489', 'Corvus brachyrhynchos', 'USA'], ['KJ501233', 'Cyanocitta cristata', 'USA'], ['KJ501320', 'Corvus brachyrhynchos', 'USA'], ['JN051152', 'Corvus', 'Mexico'], ['JN051153', 'Corvus', 'Mexico'], ['KJ501363', 'Corvus brachyrhynchos', 'USA'], ['DQ080062', 'Culicidae', 'USA'], ['KJ501276', 'Bubo virginianus', 'USA'], ['KJ501443', 'Corvus brachyrhynchos', 'USA'], ['KJ501235', 'Cyanocitta cristata', 'USA'], ['KJ501234', 'Cyanocitta cristata', 'USA'], ['KJ501282', 'Bubo virginianus', 'USA'], ['DQ080072', 'Dumetella', 'USA'], ['KJ501455', 'Cyanocitta cristata', 'USA'], ['KJ501512', 'Corvus brachyrhynchos', 'USA'], ['KX547253', 'Corvus brachyrhynchos', 'USA'], ['KJ501275', 'Corvus brachyrhynchos', 'USA'], ['DQ080071', 'Equus caballus', 'Mexico'], ['KJ501395', 'Corvus brachyrhynchos', 'USA'], ['KJ501264', 'Cyanocitta cristata', 'USA'], ['DQ431697', 'Homo sapiens', 'USA'], ['HM488236', 'Culiseta melanura', 'USA'], ['HM488218', 'Culex pipiens', 'USA'], ['DQ164202', 'Homo sapiens', 'USA'], ['GQ379156', 'Corvus', 'USA'], ['KJ501332', 'Corvus brachyrhynchos', 'USA'], ['KJ501317', 'Corvus brachyrhynchos', 'USA'], ['KJ501393', 'Corvus brachyrhynchos', 'USA'], ['KJ501516', 'Corvus brachyrhynchos', 'USA'], ['KJ501394', 'Corvus brachyrhynchos', 'USA'], ['KJ501362', 'Corvus brachyrhynchos', 'USA']]

Ecotype 57: [['KX547386', 'Culex', 'USA']]

Ecotype 58: [['HQ671708', 'Culex restuans', 'USA'], ['HM488132', 'Culiseta melanura', 'USA'], ['HQ671710', 'Culex pipiens', 'USA'], ['KX547519', 'Culex', 'USA'], ['KJ501318', 'Corvus brachyrhynchos', 'USA'], ['KJ501299', 'Corvus brachyrhynchos', 'USA'], ['KJ501494', 'Corvus brachyrhynchos', 'USA'], ['KJ501288', 'Corvus brachyrhynchos', 'USA'], ['KJ501290', 'Corvus brachyrhynchos', 'USA'], ['KJ501279', 'Cyanocitta cristata', 'USA'], ['KJ501344', 'Corvus brachyrhynchos', 'USA'], ['KJ501289', 'Corvus brachyrhynchos', 'USA'], ['KJ501343', 'Corvus brachyrhynchos', 'USA'], ['KX547478', 'Culex', 'USA'], ['KX547316', 'Culex', 'USA'], ['KX547602', 'Culex', 'USA'], ['KX547492', 'Culex', 'USA'], ['KX547228', 'Passer domesticus', 'USA'], ['KX547584', 'Corvus brachyrhynchos', 'USA'], ['KX547266', 'Culex', 'USA'], ['KX547272', 'Culex', 'USA'], ['KX547203', 'Culex', 'USA'], ['HM488136', 'Culex restuans', 'USA'], ['HM488133', 'Culex pipiens', 'USA'], ['HQ671719', 'Culex pipiens', 'USA'], ['JF920307', 'Culiseta melanura', 'USA'], ['HM488125', 'Corvus brachyrhynchos', 'USA'], ['FJ527738', 'Cyanocitta cristata', 'USA'], ['AF206518', 'Culex pipiens', 'USA'], ['HQ671706', 'Aedes vexans', 'USA'], ['HQ671707', 'Culex pipiens', 'USA'], ['JN716371', 'Phoenicopterus ruber', 'Colombia'], ['KU978766', 'Phoenicopterus ruber', 'Colombia'], ['JN716372', 'Phoenicopterus ruber', 'Colombia'], ['KX547320', 'Cyanocitta cristata', 'USA'], ['KX547564', 'Corvus brachyrhynchos', 'USA'], ['KX547395', 'Culex', 'USA'], ['HQ671714', 'Culex salinarius', 'USA'], ['HQ671697', 'Aedes vexans', 'USA'], ['FJ151394', 'Corvus', 'USA'], ['AF202541', 'Homo sapiens', 'USA']]

Ecotype 59: [['KC407667', 'Mus musculus', 'Spain'], ['AF260967', 'Equus caballus', 'USA'], ['HM488126', 'Corvus brachyrhynchos', 'USA'], ['HM488128', 'Corvus brachyrhynchos', 'USA'], ['HM488127', 'Corvus brachyrhynchos', 'USA'], ['HQ671709', 'Culex pipiens', 'USA'], ['HQ596519', 'Corvus', 'USA'], ['HQ671712', 'Culiseta melanura', 'USA'], ['DQ164194', 'Corvus brachyrhynchos', 'USA'], ['KX547353', 'Culex', 'USA'], ['AF404754', 'Culex pipiens', 'USA'], ['HM488129', 'Culex salinarius', 'USA'], ['HM152773', 'Homo sapiens', 'Israel'], ['KJ501445', 'Corvus brachyrhynchos', 'USA'], ['KJ501374', 'Buteo jamaicensis', 'USA'], ['HM488248', 'Corvus brachyrhynchos', 'USA'], ['KX547554', 'Turdus migratorius', 'USA'], ['KX547509', 'Culex', 'USA'], ['HM756661', 'Corvus brachyrhynchos', 'USA'], ['KX547179', 'Corvus brachyrhynchos', 'USA'], ['KJ501270', 'Cyanocitta cristata', 'USA'], ['KJ501375', 'Corvus brachyrhynchos', 'USA'], ['KX547538', 'Culex', 'USA'], ['DQ164188', 'Corvus brachyrhynchos', 'USA'], ['KX547194', 'Corvus brachyrhynchos', 'USA'], ['HM756663', 'Corvus brachyrhynchos', 'USA'], ['KX547261', 'Passer domesticus', 'USA'], ['HM488247', 'Corvus brachyrhynchos', 'USA'], ['HM488175', 'Culex restuans', 'USA'], ['HM756653', 'Culex pipiens', 'USA'], ['HM488246', 'Corvus brachyrhynchos', 'USA'], ['HQ671711', 'Culex pipiens', 'USA'], ['HM488130', 'Culex salinarius', 'USA'], ['KJ501515', 'Buteo jamaicensis', 'USA'], ['HM488249', 'Corvus brachyrhynchos', 'USA'], ['KJ501277', 'Corvus brachyrhynchos', 'USA'], ['KJ501278', 'Corvus brachyrhynchos', 'USA'], ['KX547251', 'Culex', 'USA'], ['KX547490', 'Corvus brachyrhynchos', 'USA'], ['KJ501378', 'Corvus brachyrhynchos', 'USA'], ['KJ501486', 'Buteo jamaicensis', 'USA'], ['KJ501450', 'Corvus brachyrhynchos', 'USA'], ['HM756662', 'Corvus brachyrhynchos', 'USA'], ['HM756664', 'Corvus brachyrhynchos', 'USA'], ['KJ501465', 'Corvus brachyrhynchos', 'USA'], ['EF530047', 'Corvus brachyrhynchos', 'USA'], ['AF404756', 'Corvus', 'USA'], ['EF657887', 'Corvus brachyrhynchos', 'USA']]

Ecotype 60: [['KX547210', 'Culex', 'USA'], ['HQ671716', 'Culiseta melanura', 'USA'], ['HQ671717', 'Aedes cinereus', 'USA'], ['HM488134', 'Ochlerotatus sollicitans', 'USA'], ['HM488135', 'Ochlerotatus cantator', 'USA'], ['HQ671696', 'Culex salinarius', 'USA'], ['KJ786934', 'Homo sapiens', 'USA'], ['AF533540', 'Homo sapiens', 'USA'], ['HQ671713', 'Corvus brachyrhynchos', 'USA'], ['HM488131', 'Culex pipiens', 'USA'], ['HQ671718', 'Culex salinarius', 'USA'], ['KX547393', 'Culex', 'USA'], ['HQ671715', 'Culex pipiens', 'USA'], ['KX547378', 'Culex', 'USA'], ['AF404753', 'Corvus', 'USA'], ['KJ501480', 'Corvus brachyrhynchos', 'USA'], ['DQ164187', 'Corvus brachyrhynchos', 'USA'], ['JN819315', 'Aves', 'USA'], ['DQ164192', 'Corvus brachyrhynchos', 'USA'], ['KJ501313', 'Turdus migratorius', 'USA']]

Ecotype 61: [['DQ164206', 'Cyanocitta cristata', 'USA'], ['AF196835', 'Gallus gallus', 'USA'], ['GQ379160', 'Equus caballus', 'Argentina'], ['GQ379161', 'Equus caballus', 'Argentina']]

Ecotype 62: [['DQ118127', 'Anser', 'Hungary']]

Ecotype 63: [['JX442279', 'Culex pipiens', 'China'], ['KC601756', 'Homo sapiens', 'India'], ['JX041634', 'Homo sapiens', 'Russia'], ['AY278441', 'Homo sapiens', 'Russia'], ['DQ374652', 'Corvus corone cornix', 'Russia'], ['DQ411029', 'Phalacrocorax carbo', 'Russia'], ['DQ411030', 'Hyalomma marginatum', 'Russia'], ['DQ411031', 'Corvus corone cornix', 'Russia'], ['DQ374650', 'Phalacrocorax carbo', 'Russia'], ['DQ374653', 'Corvus corone cornix', 'Russia'], ['DQ377178', 'Hyalomma marginatum', 'Russia'], ['DQ377180', 'Phalacrocorax carbo', 'Russia'], ['DQ411032', 'Columba livia', 'Russia'], ['DQ411033', 'Phalacrocorax carbo', 'Russia'], ['DQ411035', 'Anopheles messeae', 'Russia'], ['DQ411034', 'Anopheles messeae', 'Russia'], ['DQ374651', 'Pica pica', 'Russia'], ['DQ377179', 'Corvus frugilegus', 'Russia']]

Ecotype 64: [['KJ958922', 'Equus caballus', 'Turkey']]

Ecotype 65: [['AF260969', 'Culex pipiens', 'Romania'], ['AY278442', 'Homo sapiens', 'Russia'], ['AF317203', 'Homo sapiens', 'Russia'], ['AY277252', 'Homo sapiens', 'Russia'], ['KU588135', 'Camelus dromedarius', 'United Arab Emirates'], ['MF797870', 'Homo sapiens', 'Cyprus'], ['JQ928174', 'Homo sapiens', 'Italy'], ['JX556213', 'Homo sapiens', 'Italy'], ['KC954092', 'Homo sapiens', 'Italy'], ['KF647253', 'Homo sapiens', 'Italy'], ['AY701413', 'Equus caballus', 'Morocco'], ['DQ786572', 'Passer domesticus', 'France'], ['JF719069', 'Equus caballus', 'Spain'], ['JF719065', 'Corvidae', 'Italy'], ['JF719067', 'Laridae', 'Italy'], ['JF719066', 'Corvidae', 'Italy'], ['FJ483549', 'Passeriformes', 'Italy'], ['FJ483548', 'Passeriformes', 'Italy'], ['KF234080', 'Homo sapiens', 'Italy'], ['GU011992', 'Homo sapiens', 'Italy'], ['JF719068', 'Corvidae', 'Italy'], ['AF404757', 'Equus caballus', 'Italy'], ['FJ766332', 'Aquila chrysaetos', 'Spain'], ['FJ766331', 'Aquila chrysaetos', 'Spain'], ['JN858069', 'Homo sapiens', 'Italy'], ['JQ928175', 'Homo sapiens', 'Italy'], ['AY701412', 'Equus caballus', 'Morocco'], ['HM152775', 'Homo sapiens', 'Israel']]

Ecotype 66: [['HM051416', 'Homo sapiens', 'Israel'], ['JX041630', 'Aves', 'Azerbaijan'], ['JX041629', 'Aves', 'Azerbaijan']]

Ecotype 67: [['KX394398', 'Culex annulirostris', 'Australia'], ['KX394396', 'Culex annulirostris', 'Australia'], ['GQ851602', 'Culex annulirostris', 'Australia'], ['KX394405', 'Culex annulirostris', 'Australia'], ['KX394409', 'Culex annulirostris', 'Australia'], ['KX394383', 'Culex annulirostris', 'Australia'], ['KX394394', 'Culex annulirostris', 'Australia'], ['KT934796', 'Equus caballus', 'Australia'], ['KT934798', 'Culex annulirostris', 'Australia'], ['KX394395', 'Culex annulirostris', 'Australia'], ['KT934799', 'Culex annulirostris', 'Australia'], ['KT934803', 'Equus caballus', 'Australia'], ['KT934800', 'Culex annulirostris', 'Australia'], ['KT934801', 'Culex annulirostris', 'Australia'], ['JX123031', 'Equus caballus', 'Australia'], ['KT934804', 'Culex annulirostris', 'Australia'], ['JN887352', 'Equus caballus', 'Australia'], ['JX123030', 'Equus caballus', 'Australia'], ['KX394382', 'Culex annulirostris', 'Australia'], ['KT934802', 'Culex annulirostris', 'Australia'], ['GQ851603', 'Culex annulirostris', 'Australia'], ['KT934797', 'Homo sapiens', 'Australia'], ['KX394389', 'Culex annulirostris', 'Australia'], ['KX394390', 'Culex annulirostris', 'Australia'], ['KX394391', 'Culex annulirostris', 'Australia'], ['KX394384', 'Culex annulirostris', 'Australia'], ['KX394388', 'Culex annulirostris', 'Australia'], ['KX394387', 'Culex annulirostris', 'Australia'], ['KX394386', 'Culex annulirostris', 'Australia'], ['KX394385', 'Culex annulirostris', 'Australia']]

Ecotype 68: [['DQ256376', 'Homo sapiens', 'India'], ['KU978770', 'Homo sapiens', 'India'], ['JX041632', 'Culex vishnui', 'India'], ['GQ851605', 'Culicidae', 'India']]

Ecotype 69: [['KU978767', 'Culex quinquefasciatus', 'Madagascar'], ['FJ425721', 'Homo sapiens', 'Russia'], ['KJ934710', 'Hyalomma marginatum', 'Romania'], ['KT207791', 'Culicidae', 'Italy'], ['EF429199', 'Homo sapiens', 'South Africa'], ['KM052152', 'Equus caballus', 'South Africa'], ['EF429200', 'Homo sapiens', 'South Africa'], ['JX041631', 'Aves', 'Ukraine'], ['KC496015', 'Equus caballus', 'Hungary'], ['KJ577738', 'Homo sapiens', 'Greece'], ['KJ883346', 'Homo sapiens', 'Greece'], ['KJ883350', 'Homo sapiens', 'Greece'], ['KJ577739', 'Homo sapiens', 'Greece'], ['KF179639', 'Homo sapiens', 'Greece'], ['KJ883348', 'Homo sapiens', 'Greece'], ['KJ883344', 'Homo sapiens', 'Greece'], ['KJ883343', 'Homo sapiens', 'Greece'], ['KJ883349', 'Homo sapiens', 'Greece'], ['KJ883342', 'Homo sapiens', 'Greece'], ['KJ883345', 'Homo sapiens', 'Greece'], ['HQ537483', 'Culex pipiens', 'Greece'], ['KY594040', 'Homo sapiens', 'Greece'], ['KT757323', 'Culex pipiens', 'Serbia'], ['KT757320', 'Culex pipiens', 'Serbia'], ['KX375812', 'Homo sapiens', 'Serbia'], ['KT757322', 'Culex pipiens', 'Serbia'], ['KT757321', 'Culex pipiens', 'Serbia'], ['KT757318', 'Culex pipiens', 'Serbia'], ['KU206781', 'Homo sapiens', 'Bulgaria'], ['MH021189', 'Homo sapiens', 'Belgium'], ['KC407673', 'Accipiter gentilis', 'Serbia'], ['KF179640', 'Accipiter gentilis', 'Austria'], ['MF984345', 'Falco', 'Austria'], ['MF984351', 'Culex pipiens', 'Austria'], ['MF984338', 'Homo sapiens', 'Austria'], ['MF984352', 'Culex pipiens', 'Austria'], ['MF984349', 'Equus caballus', 'Austria'], ['MF984343', 'Homo sapiens', 'Austria'], ['MF984350', 'Equus caballus', 'Austria'], ['MF984337', 'Homo sapiens', 'Austria'], ['MF984342', 'Homo sapiens', 'Austria'], ['KP109692', 'Culex pipiens', 'Austria'], ['MF984340', 'Homo sapiens', 'Austria'], ['MF984344', 'Accipiter gentilis', 'Austria'], ['MF984346', 'Homo sapiens', 'Austria'], ['JN858070', 'Homo sapiens', 'Italy'], ['MF984341', 'Homo sapiens', 'Austria'], ['MF984348', 'Homo sapiens', 'Austria'], ['MF984347', 'Homo sapiens', 'Austria'], ['KM659876', 'Homo sapiens', 'Austria'], ['KP109691', 'Homo sapiens', 'Austria'], ['KF647252', 'Homo sapiens', 'Italy'], ['KF647249', 'Homo sapiens', 'Italy'], ['KF647251', 'Homo sapiens', 'Italy'], ['KF647250', 'Homo sapiens', 'Italy'], ['KF588365', 'Homo sapiens', 'Italy'], ['KP789960', 'Homo sapiens', 'Italy'], ['KP789958', 'Homo sapiens', 'Italy'], ['KP789955', 'Homo sapiens', 'Italy'], ['KT207792', 'Culicidae', 'Italy'], ['KP789956', 'Homo sapiens', 'Italy'], ['KP789953', 'Homo sapiens', 'Italy'], ['KP789959', 'Homo sapiens', 'Italy'], ['KF823806', 'Homo sapiens', 'Italy'], ['KP789954', 'Homo sapiens', 'Italy'], ['KP789957', 'Homo sapiens', 'Italy'], ['MF984339', 'Homo sapiens', 'Austria'], ['KC496016', 'Culex pipiens', 'Serbia'], ['KT359349', 'Homo sapiens', 'Hungary'], ['DQ116961', 'Accipiter gentilis', 'Hungary'], ['EF429198', 'Homo sapiens', 'South Africa'], ['EF429197', 'Homo sapiens', 'South Africa'], ['JN393308', 'Equus caballus', 'South Africa'], ['LC318700', 'Culex quinquefasciatus', 'Zambia'], ['KY523178', 'Culicidae', 'Uganda']]

**Supplementary Table 2. Host associations of WNV putative ecotypes with birds versus humans in the USA.** Listed are the total numbers of viruses from each ecotype infecting each host category, without regard to whether the sequences were unique. Included are those ecotypes with at least one human and one bird sample from the USA.

|  | Birds | Humans |
| --- | --- | --- |
| Ecotype 4 | 48 | 3 |
| Ecotype 11 | 1 | 1 |
| Ecotype 14 | 6 | 2 |
| Ecotype 15 | 7 | 13 |
| Ecotype 23 | 16 | 17 |
| Ecotype 24 | 12 | 2 |
| Ecotype 30 | 11 | 1 |
| Ecotype 35 | 13 | 1 |
| Ecotype 39 | 12 | 1 |
| Ecotype 40 | 47 | 2 |
| Ecotype 42 | 3 | 2 |
| Ecotype 47 | 24 | 13 |
| Ecotype 48 | 22 | 3 |
| Ecotype 50 | 19 | 1 |
| Ecotype 53 | 19 | 1 |
| Ecotype 55 | 1 | 1 |
| Ecotype 56 | 28 | 2 |
| Ecotype 58 | 16 | 1 |
| Ecotype 60 | 7 | 2 |
